# A Genetic Screen for Suppressors of Cryptic 5’ Splicing in *C. elegans* Reveals Roles for KIN17 and PRCC in Maintaining Both 5’ and 3’ Splice Site Identity

**DOI:** 10.1101/2021.06.30.450627

**Authors:** Jessie M.N.G. Lopez, Kenneth Osterhoudt, Catiana Holland Cartwright, Sol Katzman, Alan M. Zahler

**Affiliations:** Center for Molecular Biology of RNA, Department of Molecular Cell Developmental Biology, University of California, Santa Cruz, Santa Cruz, California, United States of America; Center for Biomolecular Science and Engineering, University of California, Santa Cruz, Santa Cruz, California, United States of America

## Abstract

Pre-mRNA splicing is an essential step of eukaryotic gene expression carried out by a series of dynamic macromolecular protein/RNA complexes, known collectively and individually as the spliceosome. This series of spliceosomal complexes define, assemble on, and catalyze the removal of introns. Molecular model snapshots of intermediates in the process have been created from cryo-EM data, however, many aspects of the dynamic changes that occur in the spliceosome are not fully understood. *Caenorhabditis elegans* follow the GU-AG rule of splicing, with almost all introns beginning with 5’ GU and ending with 3’ AG. These splice sites are identified early in the splicing cycle, but as the cycle progresses and “custody” of the pre-mRNA splice sites is passed from factor to factor as the catalytic site is built, the mechanism by which splice site identity is maintained or re-established through these dynamic changes is unclear. We performed a genetic screen in *C. elegans* for factors that are capable of changing 5’ splice site choice. We report that KIN17 and PRCC are involved in splice site choice, the first functional splicing role proposed for either of these proteins. Previously identified suppressors of cryptic 5’ splicing promote distal cryptic GU splice sites, however, mutations in KIN17 and PRCC instead promote usage of an unusual proximal 5’ splice site which defines an intron beginning with UU, separated by 1nt from a GU donor. We performed high-throughput mRNA sequencing analysis and found that mutations in PRCC but not KIN17 changed 5’ splice sites genome-wide, promoting usage of nearby non-consensus sites. We further found that mutations in KIN17 and PRCC changed dozens of 3’ splice sites, promoting non-consensus sites upstream of canonical splice sites. Our work has uncovered both fine and coarse mechanisms by which the spliceosome maintains splice site identity during the complex assembly process.

**Author Summary:** Pre-mRNA splicing is an essential step of gene regulation, carried out by an unusual molecular machine, the spliceosome. Unlike other molecular machines, such as ribosomes, that simply assemble and catalyze chemical reactions, “the spliceosome” is a highly-dynamic cycle, carried out by 5 specialized small nuclear RNAs and over 100 proteins, which sequentially join, rearrange, and withdraw from the splicing assembly during each splicing cycle. These assemblies initially choose “splice sites” where the pre-mRNA will be cut, and then undergo multiple rearrangements to finally form the active site which catalyzes the splicing reactions which remove an intron from a pre-mRNA. We are currently in the midst of a “resolution revolution”, with ever-clearer cryo-EM snapshots of stalled complexes allowing researchers to visualize moments in time in the splicing cycle. These models are illuminating, but do not always elucidate mechanistic functioning, therefore our lab takes a complementary approach, using the power of genetics in a multicellular animal to gain functional insights into the spliceosome. Using a *C .elegans* genetic screen, we have found novel functional splicing roles for two proteins, KIN17 and PRCC. Our results suggest that the spliceosome does not just rely on its initial identification of the splice site, but in a later step, re-identifies where to cut. We liken this two-stage identification to using a microscope by first using the coarse focus to find the area of interest, and then using the fine focus to adjust as needed. This work moves us closer to full mechanistic understanding of how the spliceosome chooses where to cut a pre-mRNA message.

## Introduction

The spliceosome is not one distinct machine but a series of dynamic macromolecular protein/RNA complexes that assemble on and catalyze the removal of introns from pre-mRNA transcripts in eukaryotic organisms. Over one hundred proteins, including multiple helicases, and the 5 U-rich small nuclear RNAs (snRNAs) join, rearrange, and withdraw from a spliceosomal complex in a choreographed sequence over the course of a single splicing cycle, catalyzing the removal of an intron, and ligation of the flanking exons [1,2]. Spliceosomes assemble *de novo* from subunits on each nascent pre-mRNA intron, Multiple spliceosomes often interact with a pre-mRNA transcript at the same time, and different introns in a pre-mRNA can have different kinetics for removal [3]. The splicing process is responsible for an essential information processing step in the flow of genetic information, and almost all protein-coding transcripts in metazoans must be spliced in order to become functional.

Early in the metazoan splicing cycle, three important landmarks on the nascent pre-mRNA are identified by spliceosomal components: the 5’ splice site (exon/intron boundary), the branchpoint, and the 3’ splice site (intron/exon boundary). The U1 snRNA has a 9 base sequence, 3’ GUCCAψψCAUA 5’ that pairs with the bases of the 5’ splice site [4]. A perfectly complementary 5’ splice site would have the sequence 5’ CAG/GUAAGUAU 3’, where the slash represents the splice site, however this exact sequence is rarely found at verified 5’ splice sites in metazoans. Instead, a consensus sequence that has some overall base pairing ability with U1snRNA, with a strong preference for a /GU dinucleotide to start the intron, is seen [5]. The /G is nearly invariant, its 5’ phosphate will link directly to the branchpoint adenosine. For the 3’ss, the U2AF heterodimer initially identifies the polypyrimidine tract and AG dinucleotide at the end of the intron; U2AF65 binds the polypyrimidine tract, and U2AF35 binds the nearly invariant AG/ at the very 3’ end of the intron [6]. U2AF helps to recruit U2 snRNP to the branch site where base-pairing interactions with U2snRNA, in which the branch point adenosine is bulged out of the duplexed region, define the branchpoint [6,7].

Mutations in splice sites or in cis-regulatory regions, such as enhancer or silencer binding sites, can cause a variety of deleterious splicing phenotypes that are associated with disease phenotypes. Examples include exon skipping, intron inclusion, and frameshift mutations. Mutation of a splicing donor or acceptor sequence leads to activation of nearby “cryptic” splice sites, which are defined as splice sites that are functional but activated only when an authentic splice site is disrupted by mutation. In the Human Gene Mutation Database, ~9% of inherited disease-causing mutations alter splice site sequences [8], and another ~25% of disease-causing mutations affect splicing by disrupting other important sequences, such as nearby binding sites [9,10]. Some aberrant mRNAs are degraded by non-stop, or nonsense-mediated decay pathways, so that the possibly toxic effects of aberrant mRNAs are not amplified into many aberrant proteins by polyribosomes [11]. Precise splicing is central to gene expression, and mutations that affect splicing can lead to a variety of deleterious phenotypes.

Throughout the many dynamic assembly steps of the splicing cycle, the U1-identified 5’ splice site is maintained by a series of protein and snRNA escorts. In the earliest steps of spliceosome assembly, the 5’ splice site is directly bound by U1 snRNA [12]. In the transition from pre-B to B-complex, U1 leaves the spliceosome while handing the 5’ splice site off to U6 and residues of PRP8 [13,14]. From B complex, the spliceosome undergoes a number of rearrangements through pre-Bact1, pre-Bact2, Bact and C complex. CryoEM studies of these complexes from human spliceosomes [2,15] allow for the study of different snapshots of the spliceosome assembly process. In these complexes there is an exchange of different factors that interact with the region of the 5’ss and its interactions with the U6 ACAGAGA box as the 5’ss is loaded into the catalytic core of the splicing machine. Proteins and snRNPs that bind to the 5’ splice site must bind precisely to a degenerate sequence on a long nucleotide chain, maintain their exact binding position through helicase-powered translocations and substantial conformational changes, and then transfer custody of the 5’ splice site to the next escort, without introducing positional error. It is still unclear which components of the spliceosome ensure that the handoffs between escorts will not result in small shifts in 5’ splice site definition.

Thanks to the researchers fueling the ongoing cryo-EM resolution revolution, we now have structures of spliceosomes at many time points in the splicing cycle. These snapshots of experimentally stalled spliceosome assemblies offer valuable insights into the complex assembly pathways, rearrangements and interactions of spliceosomal components [2]. Mass spectrometry experiments and chemical probing of structures have provided additional information about where and when specific components are associated with the spliceosome during the splicing cycle. These advances continue to build towards a fuller picture of the many multi-step assembly pathways of the splicing cycle and the organized dissolution of the complex. While the structuralists reveal which proteins are where, geneticists are positioned to provide complementary insights into the functional roles of splicing components in splice site choice.

Our lab has previously made use of an unusual 5’ splice site mutation in *C. elegans* as a tool to reveal residues on splicing proteins that can contribute to splice site choice [16] [17]. UNC-73 is a guanine nucleotide exchange factor that is important in axon guidance and other aspects of *C. elegans* development. A fortuitous G->U mutation of the first nucleotide of the 16th intron of the *unc-73* gene, allele *e936* (ce10::chrI:4,021,954) [18] converts the nearly invariant /GU dinucleotide found at the beginning of eukaryotic introns to a /UU dinucleotide, creating a curiously ambiguous splice site (Fig 1A). This splice site mutation results in misplicing, causing the uncoordinated (unc) phenotype [19]. This dramatic phenotype is corrected by even a small increase in in-frame splicing, making its suppression screenable. Previously identified dominant mutations that are able to suppress the unc phenotype by altering cryptic splicing in *unc-73(e936)* were found in U1snRNA [20],] SNRP-27 [16]; [21] and the largest and most conserved protein in the spliceosome, PRP-8 [17]. The suppressive role these mutations play in this splice site assay provided genetic evidence of a role for these protein residues in 5’ splice site choice. After publishing these data, the progress made in cryo-EM and crystal structures of the spliceosome has allowed these suppressor alleles to be precisely mapped in the high-resolution inner core of spliceosomal structures; these mutations are often modeled near the active site of the spliceosome providing some clues as to mechanisms for maintaining the identity of the 5’ss during spliceosome assembly.

**Figure 1.**
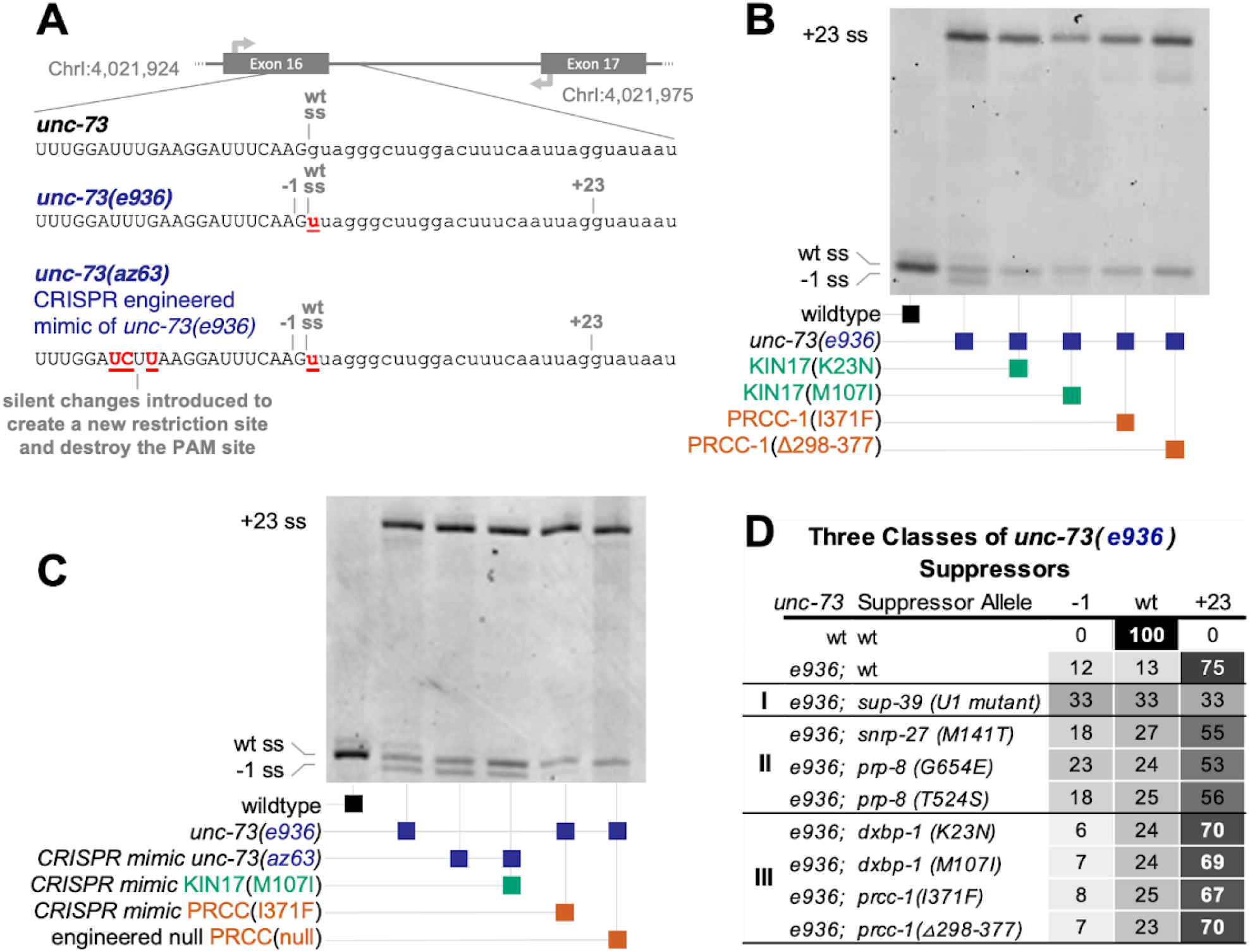
Mutations in KIN17/dxbp-1 and PRCC suppress cryptic splicing, promoting an unusual /UU 5’ splice site. **(A)** Schematic diagram of the 16th intron of the C. *elegans* gene *unc-73*, showing genomic coordinates and relative loci of splice sites and PCR primer locations used to assess splice site usage. Below, aligned sequences of the *unc-73* sequence and exon/intron boundary in wild type, *unc-73(e936)*, and in the CRISPR engineered allele *unc-73(az63).* The cryptic splice sites activated in the competition assay are labeled −1 and +23 and define introns beginning with /GU that are both out of frame. Note that the wild-type splicing position is still denoted “wt ss” even though that intron now begins /UU. The slash mark (/) denotes the splice site. **(B)** Poly-acrylamide gel showing Cy-3 labeled *unc-73* PCR products amplified from *unc-73* cDNA. RNA was extracted from plates of the following 6 strains of C. *elegans*: wild-type N2, *unc-73(e936)*, and four independent original suppressed strains identified in the genetic screen whose genotypes are indicated below, each bears both the unc-73(e936) allele and an extragenic suppressor of both the movement defect. The same PCR primers are used on all samples; band positions and intensities are indicative of relative use of the three available 5’ splice sites, labeled −1, wt, and +23. Strains are, in lane order, N2, SZ181, SZ162, SZ283, SZ280, and SZ281, see Methods for genetic details. **(C)** Putative suppressor identities were verified by *de novo* recreations of mutations using CRISPR/Cas9 and homology-directed repair into *unc-73* reporter strains. Image is a scan of a denaturing poly-acrylamide gel showing Cy-3 labeled *unc-73* PCR products from *unc-73* cDNA. RNA was extracted from strains with the indicated *unc-73, dxbp-1*, and *prcc-1* alleles shown below. Strains are, in lane order, N2, SZ181, SZ219, SZ222, SZ308, SZ348, see Methods for genetic details. Unless otherwise mentioned, CRISPR-engineered mimic alleles are used for all subsequent experiments and figures in this report. **(D)** Four new suppressors of cryptic splicing represent a new class of suppressors, with a distinct molecular phenotype compared to previously identified suppressors. Table rows show suppressor class (I, II, or III) [16,17,20-22], genotype of *unc-73*, genotype of suppressor, average percent splice in (PSI), n>3, at the /GU splice site at position −1 relative to wild type, average PSI at /UU splice site in wild-type position, and average PSI at the /GU at position +23. Conditional grayscale shading highlights patterns in numerical data.

Here we report new additional suppressor alleles identified in the *unc-73(e936)* genetic screen for suppression of uncoordination that have a dramatically different mechanism of suppression through splicing. Previous suppressors promoted the use of both the −1 and wt cryptic sites separated by 1nt, /G/UU, over a downstream cryptic GU splice donor at position +23. Here we identify two new proteins as splicing factors in which mutations promote use of the /UU splice donor over the adjacent GU splice site. Two missense alleles in the worm homolog of **KIN17** (Kinship to RecA), called *dxbp-1* (downstream of x-box) in *C. elegans*, and an overlapping point mutation and deletion in the worm homolog of human **PRCC** (proline-rich coiled coil protein or papillary renal cell carcinoma protein), called *prcc-1* in *C. elegans*, promote the usage of an unusual /UU splice site in 3-choice, 2-choice and 2X2-choice cryptic splice site assays. High throughput mRNA-SEQ studies reveal that these mutations affect global splicing at native splice sites, but despite similarities in effects on *unc-73(e936)* cryptic splicing, mutations in KIN17 and PRCC display strong, but very different, effects on native genes. These results are the first demonstration that PRCC and KIN17 have roles in maintaining splice site identity during spliceosome assembly.

## Results

### Mutations in KIN17 and PRCC alter cryptic 5’ splice site choice in *unc-73(e936)*

The *unc-73(e936)* allele has a G→U mutation at the 1st nucleotide (+1) position of the 16th intron. This mutation presents the spliceosome with an ambiguous 5’ splice site, resulting in the usage of two out-of-frame cryptic 5’ss and a striking uncoordinated phenotype [19] (Figure 1A). The majority of splicing (75%) occurs at a /GU dinucleotide found 23 nucleotides into the intron (the +23 site), resulting in an out-of-frame message. An additional 12% of splicing occurs at a position 1nt upstream of the wild-type splice site (the −1 site) using the new /GU dinucleotide formed by the *e936* mutation, also resulting in an out-of-frame message. These out-of-frame messages are not substrates for nonsense-mediated decay [19]. An additional 13% of splicing occurs at the wild-type splice site (the wt site), even though this defines an intron that begins with a non-canonical /UU. Only the small fraction of splicing at the in-frame /UU splice site produces full-length functional protein. The animals bearing the *unc-73(e936)* allele are able to live and reproduce through self-fertilization, but are profoundly uncoordinated. Even a modest increase in splicing at the in-frame /UU splice site results in a dramatic phenotypic reversal which is visible at the plate level, making this allele a sensitive assay of perturbations to splice site choice [16,17,19]). Because those previous screens have identified mutations on residues modelled near the active site of the spliceosome, and those mutations often change global 5’ splice site choice, we concluded that a genetic screen using this allele can identify loci which are capable of affecting splice site choice. Because we have never found the same mutation twice in 500,000 mutagenized genomes screened previously, we hypothesized the screen is not yet saturated. Therefore, we performed the genetic screen again to search for more suppressor mutations in splicing factors capable of altering splice site choice.

In a recent iteration of the *e936* extragenic suppressor screen, we recovered four new extragenic suppressor alleles with improved locomotion and a novel change in cryptic splicing. Using Cy-3 labeled primers in reverse transcription - polymerase chain reaction (RT-PCR) visualized after denaturing gel electrophoresis, we found that these four strains displayed a different pattern of cryptic 5’ splice site usage in *unc-73* compared to wild type, but, curiously, also a different pattern compared to previously identified modifiers [16,17,19]. While previous suppressors have reduced splicing at the +23 splice site with coordinated gains at both the −1 and wt sites, these four new suppressors had the most dramatic effect in altering the relative usage of the −1 and wt sites relative to each other, resulting in increased wt splice site usage to ~25% of *unc-73* messages, consistent with the improved locomotion phenotype identified in the screen (Fig. 1B). We now refer to extragenic suppressors in three classes: Type I is the U1 snRNA suppressor *sup-39*, while Type II includes the protein factor suppressor alleles *snrp-27* (M141T) and *prp-8* T524S and G654E. The Type I and Type II suppressors both reduce +23 splice donor usage with concomitant increases in both the −1 and wt splice sites. The dramatic change in the relative −1 and wt usage is the key feature of these new Type III suppressors.

### The four new type III suppressor alleles capable are in the C. elegans homologs of KIN17 and PRCC

Using Hawaiian strain SNP mapping [26], as described in Methods, we mapped each of these four new suppressor alleles to an arm of a chromosome. Then, using high throughput DNA sequencing of the strain genomes, followed by SNP identification protocols to identify differences in genomic sequence from the starting *unc-73(e936)* uncoordinated strain (see Methods), we identified spliceosome-associated proteins and RNA binding proteins with mutations in their sequence within the chromosomal delimited chromosomal region.

Two of the suppressor alleles had point mutations in the gene *dxbp-1*, the worm homologue of KIN17: a mutation that changes the 23rd amino acid from a lysine to an arginine (K23N, *az105*, Fig 1B, Lane 3) and another that changes the 107th amino acid from a methionine to an isoleucine (M107I) (*az33*, Fig 1B, Lane 4). Both of these residues are conserved between worm, human, yeast and arabidopsis (Fig 2). *C. elegans dxbp-1*, downstream of x-box binding protein, or *dox-1*, is the homolog of a human and mouse gene known as KIN or KIN17. It is *not* a kinase. Except in the multiple sequence alignment (Fig 2), throughout this manuscript we will refer to KIN17 when talking about the protein, and *dxbp-1* when talking about the gene. The 23rd residue of the worm homolog of KIN17 is proximal to the CHC2 zinc finger in a region predicted to be near the U6 pre-mRNA helix in B^act^2 [15,24] (Fig 3). The 107th residue of the worm homolog of KIN17 resides in a 3_10_ helix on a loop in the atypical winged helix domain. This domain is atypical because the cluster of residues that are typically positively charged and coordinate nucleic acid binding in a winged helix is not charged, leading to the hypothesis that the highly conserved 3_10_ helix is involved in protein binding [24]. KIN17 is predicted to have a disordered central region flanked by α-helices [15], followed by a tandem of SH3-like domains separated by a flexible linker.

**Figure 2.**
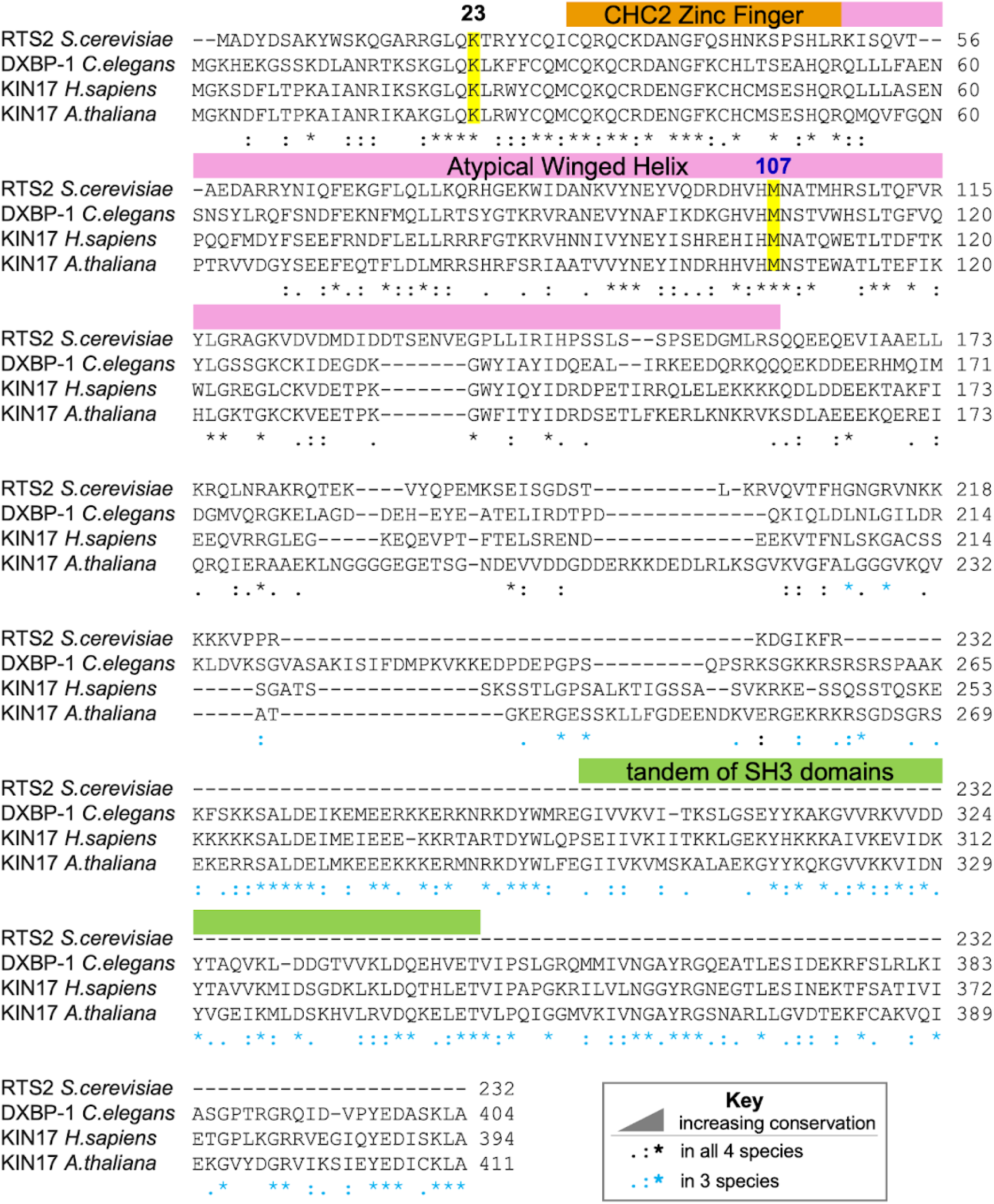
N-terminus of KIN17 is Highly Conserved Between Yeast, Worm, Human and Arabidopsis. Multiple sequence alignment of KIN17 and orthologs. K23 and M107 are highlighted in yellow, the region of the zinc finger indicated in orange, the atypical winded helix in blue, and the tandem of SH3 domains in green. Sequence conservation is annotated as described in the key. Alignment generated in Clustal Omega [23].

**Figure 3.**
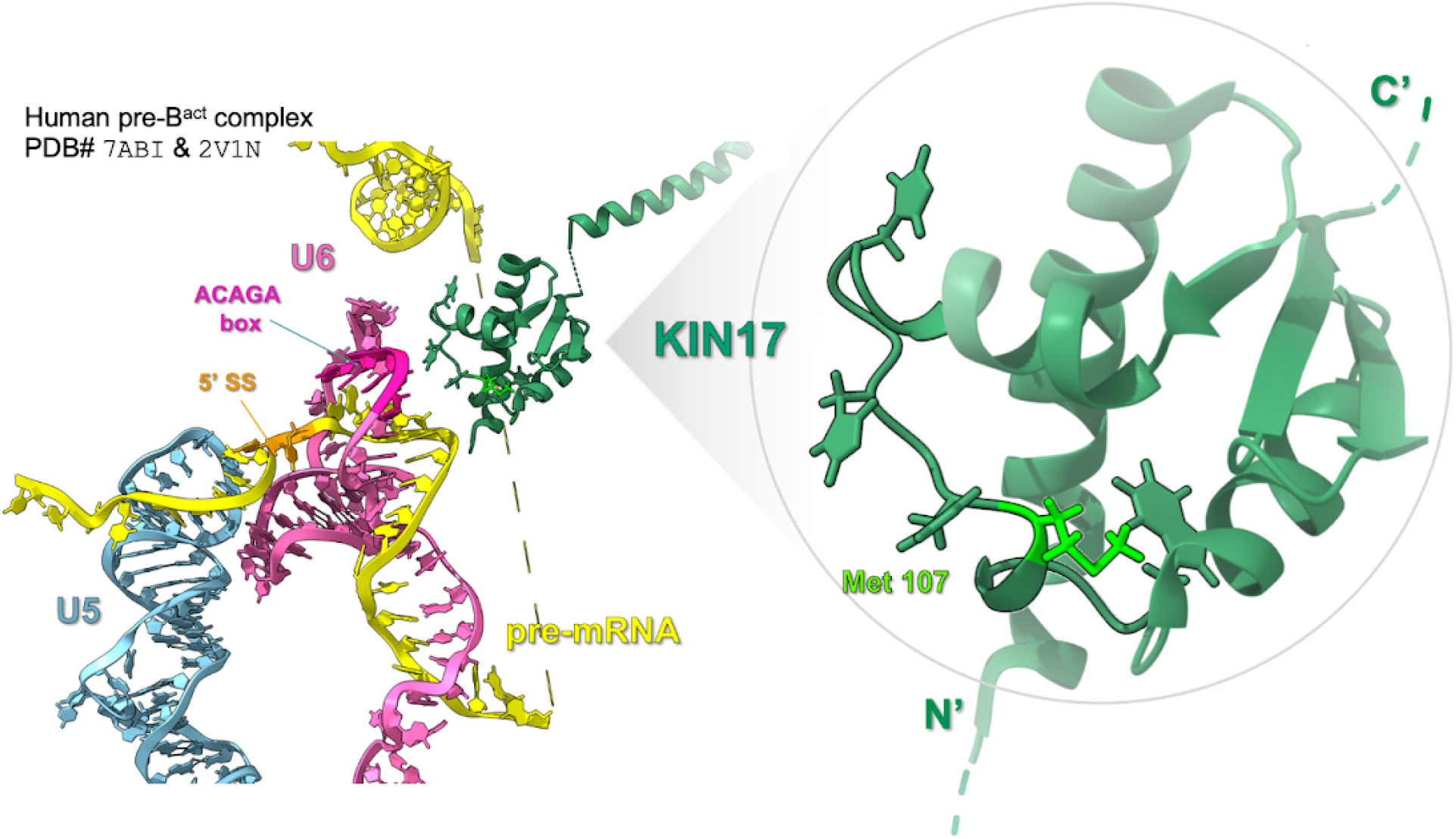
KIN17(K23N) and KIN17(M107I) are close to the pre-mRNA in human pre-B^act^2. Main figure shows a model of human U6 (pink), U5 (light blue) and the pre-mRNA (yellow) in pre-B^act^2 complex of the splicing cycle, based on Protein Data Bank structures 7ABI [15] and 2V1N [24], KIN17 (green) is near the U6/pre-mRNA loop. The pre-mRNA intron is unstructured behind KIN 17. KIN17 is magnified in the inset circle, methionine 107 (spring green) is part of a short 3_10_ helix, on a loop between two alpha helices of the winged helix. Methionine 107 is 15 A from the mRNA, and the residue points into the globular core of the winged helix domain. The 23rd residue of KIN17 is not modelled in this structure, however the N terminus of KIN 17, containing the zinc-finger and the unstructured region, points down and back into the same region as the unstructured pre-mRNA passing behind KIN 17.

KIN17 was first identified in a search for mammalian homologs of the bacterial DNA repair protein RecA, and has since been studied primarily for roles in DNA damage repair and transcription in eukaryotic cells [27–37] or cancer [38,39]. In *S. cerevisiae*, there is a named gene, RTS2, that shares homology with the N-terminal portion of KIN17 [40]. Observations about KIN17 include the following: KIN17 binds to single-stranded and double-stranded DNA [37,41–45] with a preference for AT-rich curved double-stranded DNA [31,46,47] and binds to RNA, with domains exhibiting preferences for specific poly-nucleic acid oligos [48,49]. KIN17 also binds to proteins in complexes of high molecular weight, including ones involving chromatin [41,45,50], DNA recombination [46], DNA damage repair [51], DNA replication [36,44], pre-mRNA splicing [48,52–55] [15], and translation [45]. It is likely that KIN17 performs more than one role in the eukaryotic cell.

This screen also identified two mutations in *prcc-1*, the worm homolog of human PRCC: a mutation which changes the 371st amino acid from an isoleucine to a phenylalanine (I371F) (*az102* Fig 4), and a large deletion near the C terminus that removes amino acids 298-377 in frame (*az103*, Fig 4). Except in the multiple sequence alignment, throughout this manuscript we will refer to PRCC when talking about the protein and *prcc-1* when talking about the gene. PRCC, known variously as proline-rich protein, proline-rich coiled coil, papillary renal cell carcinoma translocation-associated gene protein, and mitotic checkpoint factor protein, has been implicated in oncogenic fusions where the proline-rich N terminal region is fused to any of several transcription factors [56–58]. The proline-rich region is relatively proline-poor in *C. elegans* compared to human; the domain is absent in arabidopsis. The 371st amino acid of the worm homolog of PRCC occurs in the middle of the longest stretch of identity, where 9 residues are conserved from worm to human. The deletion suppressor identified in this screen overlays that region. (Fig 4). PRCC has been identified as a potential spliceosomal B^act^ complex component by mass spectrometry [59] and Yeast 2 Hybrid [60].

**Figure 4.**
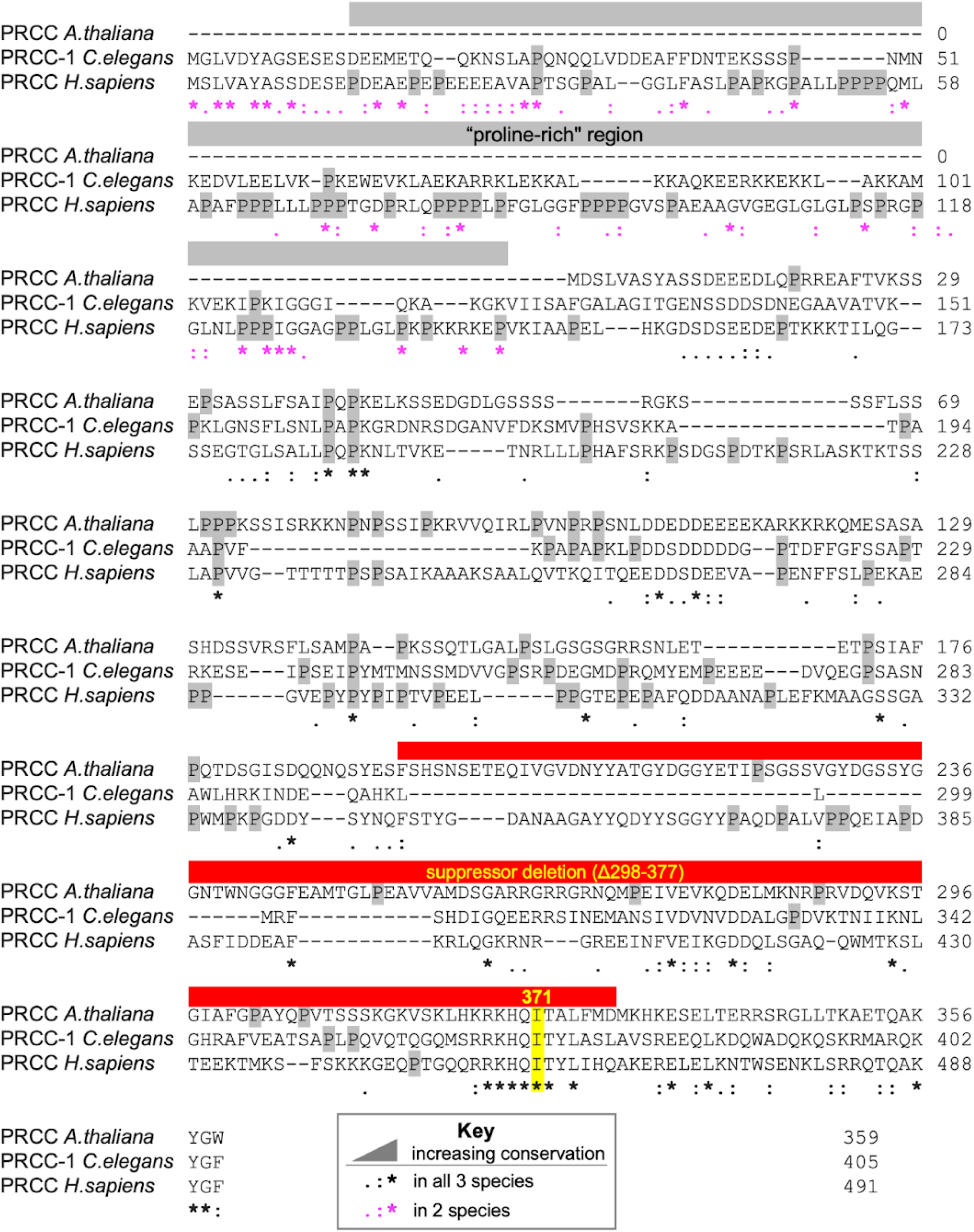
Both suppressor mutations overlap with the longest stretch of identity in PRCC. Multiple sequence alignment of PRCC-1 and orthologs. The “proline-rich” region frequently observed in oncogenic fusions is indicated in gray, and all prolines are highlighted in gray, the suppressor mutation 1371 is highlighted in yellow, the suppressor deletion (Δ289-377) is indicated in red. Sequence conservation is annotated as described in the key. PRCC(null) is not represented because it is a deletion of all coding regions of the gene. Alignment generated in Clustal Omega [23].

To confirm that the three single amino acid substitution alleles are indeed responsible for the altered cryptic splicing of *unc-73(e936)*, we used CRISPR/Cas9 to generate the same amino acid substitutions from scratch (see methods) and tested these programmed alleles for an effect on the ratio of −1:wt splice site usage. The CRISPR-generated *prcc-1(az102)* allele can suppress *unc-73(e936)* splicing and movement defects, confirming the identity of the PRCC(I371F) suppressor (Fig 1C, Lane 5). A deletion null allele of *prcc-1* generated by the *C. elegans* gene knockout consortium, *gk5556*, is viable and can both suppress the movement defects of *unc-73(e936)* and alter cryptic splice site usage (Fig 1C, Lane 6). This demonstrates that *prcc-1* is a non-essential gene and that loss-of-function leads to changes in splicing.

Confirmation of the *dxbp-1* alleles by CRISPR is a little more challenging, as they map to the same chromosome as *unc-73*, making crosses difficult, and injection of CRISPR-cas9 RNP complexes into *e936* animals is challenging as the worms are sick and have smaller brood size. We solved this challenge by generating the two *dxbp-1* mutation alleles by CRISPR in a wild-type strain, followed by subsequent CRISPR mutation of *unc-73* to mimic the *e936* allele. These strains resulted in suppression of *unc-73* uncoordination and the predicted change in −1:wt splice site usage (Fig 1C, Lanes 3 and 4). To understand whether KIN17 is an essential gene, we used our standard CRISPR pipeline to generate a *dxbp-1(null))* allele (see methods). We put the *dxbp-1(null)* allele over a fluorescent hT2 balancer, designed such that homozygous *dxbp-1(+)* animals are GFP+ but homozygous lethal, heterozygous animals are GFP+, and animals homozygous *for dxpb-1(null))* do not fluoresce. We found that KIN17 deletion is embryonic lethal in *C. elegans*; occasionally GFP- animals homozygous for *dxbp-1(null))* can survive to something resembling L3 stage, however these rare animals are severely underdeveloped and do not live to molt again. Simultaneously, the *C. elegans* Deletion Mutant Consortium [61] created a *dxpb-1(null))* allele and also found the deletion of *dxbp-1* to be homozygous lethal.

### KIN17 and PRCC promote usage of a non-canonical /UU 5’ splice site in 2-choice and 2×2-choice reporters

We were interested in the unique suppressive phenotype displayed by the mutations in KIN17 and PRCC, so similar to each other but distinct from previously identified suppressor phenotypes in that they change the relative 5’ss usage of overlapping /G/UU splice sites. In order to investigate this further, we utilized an intragenic suppressor allele of *unc-73, e936az30*, in which an A→G mutation at the +26 position of the intron eliminates the usage of the +23 cryptic splice site (Fig 5A). Therefore, the only two splice sites available are the cryptic /GU and the non-canonical /UU one nucleotide downstream; we refer to it as a 2-choice splice substrate. In a wild-type background, these two splice sites are used about 41% and 59% of the time, respectively. (Fig 5B, Lane 3)

**Figure 5.**
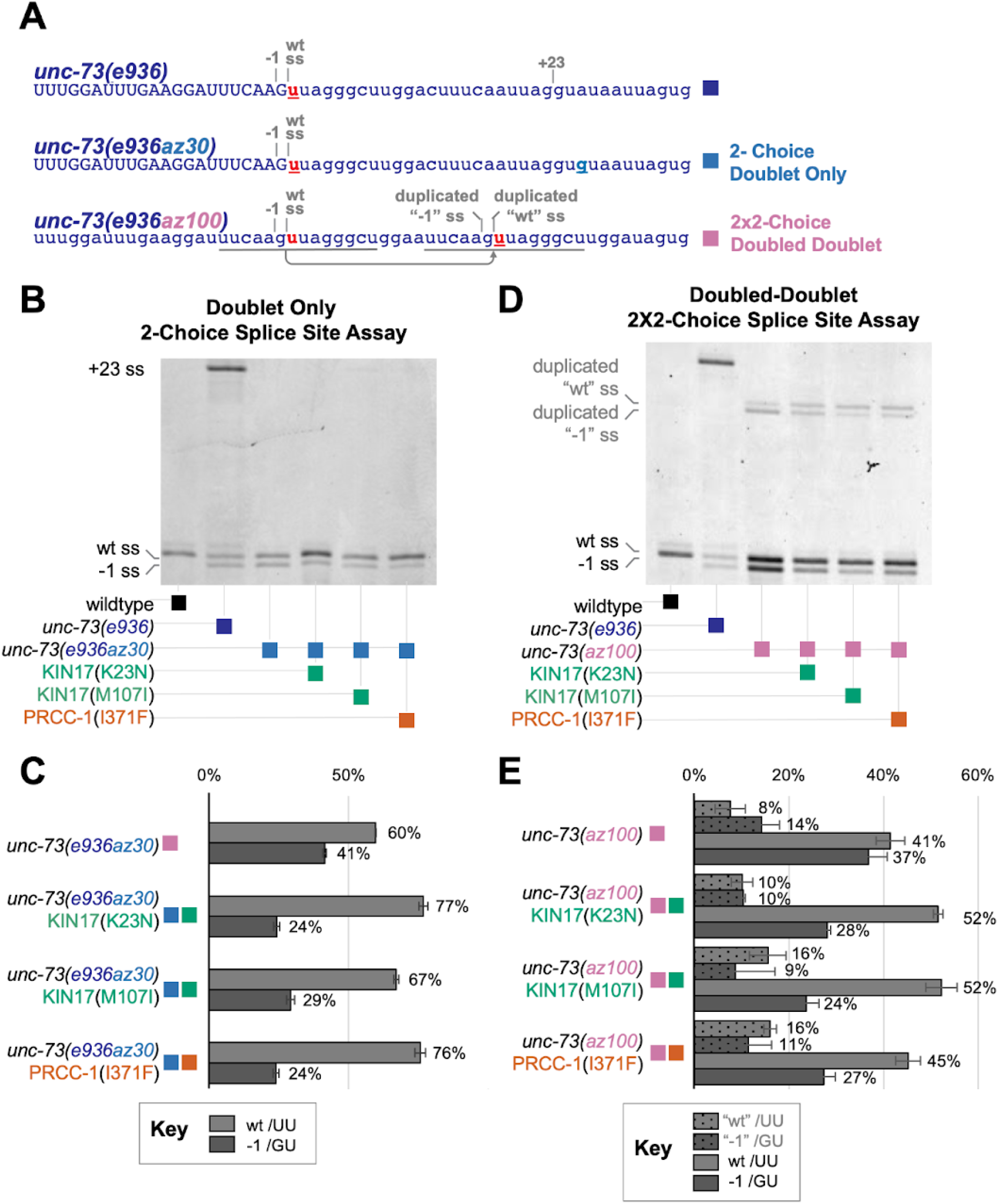
UU/ preference is independent of splice site location. **(A)** Sequences of three splice site choice competition reporters based on C. *elegans unc-73\* the first is the *unc-73(e936)* allele that allows for three cryptic splice sites as described in Figure 1A below that, *unc-73(**e936az30***) intragenic suppressor allele in which the +23 splice site is abolished by a A→G at the +26 position of the intron, leaving only the doublet of/G/UU splices sites, which we refer to as the 2-choice doublet-only splicing assay, and *unc-73(**az100**)* in which the genomic region of the doublet splice site has been duplicated, overwriting the downstream wild-type sequence and creating two /G/UU doublets, 18 bases away from each other, which we refer to as the 2×2 doubled-doublet splicing assay. **(B)** All three suppressors change the ratio of splice site usage at the doublet, promoting the /UU splice site. Poly-acrylamide gel showing Cy-3 labeled *unc-73* PCR products from cDNA. The alleles found in each sample are indicated in the figure.The same PCR primers are used on all samples; band positions and intensities are indicative of relative use of the available 5’ splice sites. **(C)** Quantification of PSI of the indicated strains, n⚕3 per strain. Error bars show Standard Deviation. **(D)** All three suppressors change the ratios of splice site usage at both the original doublet and the duplicated doublet, promoting the /UU splice site. Poly-acrylamide gel showing *unc-73* Cy-3-labeled PCR products from cDNA from the indicated strains with the indicated alleles. **(E)** Quantification of PSI of the indicated strains; details in Methods. Error bars show Standard Deviation.

In a KIN17(K23N), KIN17(M107I) or PRCC(I371F) background, we see altered ratios of splice site use in the 2-Choice splice site competition assay relative to wild-type background (Fig 5B). The splicing pattern was similar in the presence or absence of the + 23 /GU splice site (compare with Fig 1C). Despite the /GU being the primary hallmark of the 5’ splicing landmark, these suppressor alleles are promoting usage of the adjacent /UU 5’ss. In the KIN17(K23N), KIN17(M107I) and PRCC(I371F) strains, the relative /UU splice site usage is increased to 77%, 67% and 76%, respectively (Figs 5B and 5C).

In the 2-Choice splice site competition assay, we found that mutations in PRCC and KIN17 promote usage of a non-canonical /UU splice donor over an adjacent upstream /GU splice site. We wondered whether the information to promote /UU splicing was contained within the 5’ss itself, whether it was promoted by some nearby splicing enhancer element, or whether it was dependent on a distance from the original splice site. To answer these questions, we devised a new competition assay which would separate sequence from location. Using CRISPR/Cas9 and a repair oligo, the region bearing the curious /G/UU 5’ss doublet was duplicated in the native *unc-73* gene, and inserted downstream, overwriting the downstream bases of the intron (Fig 5A, allele *az100*). This doubled the splice donor doublet, creating a 2×2-choice splice site assay, featuring two 2-choice splice site doublets, 18 bases away from each other. We knew the second doublet was close enough to be chosen by the spliceosome because it was proximal to the + 23 site from the 3-choice splice site assay in the original *unc-73(e936)* allele. We abolished the + 23 splice site, so that only the four choices contained in the two doublets remained. In a wild-type background, both splice sites of the original doublet are used more than either of the splice sites in the duplicated doublet downstream. In the upstream doublet, there is a slight preference for the /UU splice site (53%), while in the less-used downstream doublet the /UU site is less-preferred (34%). (Fig 5D, Lane 3).

When this “doubled-doublet” *unc-73(az100)* allele is combined with suppressor alleles KIN17(K23N), KIN17(M107I) or PRCC(I371F), we see altered ratios of splice site use in the 2×2-Choice splice site competition assay relative to wild type (Fig 5D). In all three cases, both doublets are used and most splicing comes from the upstream doublet. In the presence of any of these three suppressor alleles, the usage of the /UU splice site increases relative to the /GU splice site in both the original doublet and the duplicated doublet, 18 nucleotides downstream. When the ratio of splice site usage at each doublet is considered independently, for KIN17(K23N), KIN17(M107I) and PRCC(I371F) we see that for both doublets, usage of the /UU splice site is increased (Fig 5E). These data support the hypothesis that the information for switching to /UU splice donor usage in the presence of these suppressor alleles is dependent on the 5’ss sequence and not a distance from some other markers on the pre-mRNA.

### Analysis of splicing changes in native genes in the presence of KIN17 and PRCC suppressor alleles

Because mutations in KIN17 and PRCC are able to promote usage of 5’ /UU splice sites in our splice site competition assays, we wanted to know if those mutations also changed splice site choice at native loci. The *unc-73* transcript, upon which all of our splice site competition assays are built, is not subject to nonsense-mediated decay [19], which is why we are able to recover cryptically-spliced frame-shifted transcripts. However, when looking for alterations displaying site choice more broadly, we expect that most transcripts will be targeted by nonsense mediated decay (NMD), especially given that the prominent splicing change we might expect to see would move the start site of an intron over by a single nucleotide, thus changing the reading frame. Given that many out-of-frame messages would be targeted by NMD, it might be difficult to detect these changes in splicing as they may potentially lead to differential transcript stability. *C. elegans* is a rare metazoan able to survive without a functional NMD pathway, making it possible to do the experiment in an NMD knockout background [62]. We designed a CRISPR/Cas9 engineered *smg-4* null allele, *az152*, which is easily detectable by single worm PCR and restriction digest, allowing for ease of mapping in crosses; *smg-4* was chosen for creating an NMD mutant strain as it is not located on the same chromosome as *dxbp-1* or *prcc-1*. We confirmed that the new *smg-4* allele is NMD-defective by both the presence of the protruding vulva phenotype and the accumulation of NMD-targeted isoforms of *rpl-12* (data not shown) [63].

We used genetic crosses to create strains with KIN17(K23N), KIN17(M107I), PRCC(I371F) or PRCC(null) combined with *smg-4(az152)*, isolated mRNA and performed mRNA-seq on three biological replicates for each suppressor strain, as well as on the original *smg-4(az152)* mutant strain as a control, 15 libraries in total. We performed 75×75bp paired end reads and obtained between 46M and 69M reads for each library. We performed star mapping, which we modified to accommodate /UU 5′ splice sites as described in Methods. We ran an alternative splicing analysis which looked at both annotated and unannotated alternative 5’ and 3’ splicing events, as well as Ensembl-annotated skipped exon, mutually exclusive exon, multiply skipped exon, intron inclusion, alternative first exon and alternative last exon events. For each alternative splicing event, we quantified relative usage of each junction in each of the 15 libraries. We then compared the percent spliced in (PSI) for each event between each library and the starting *smg-4* mutant strain. We performed pairwise comparisons between each of the three biological replicates of a suppressor strain against each of the three biological replicants of the NMD mutant strain alone, for a total of 9 comparisons for each alternative splicing event, and asked how many of those 9 comparisons generated a delta PSI of >15%. Those events for which all 9 pairwise comparisons had a delta PSI >15% (pairSum=9) were then analyzed individually on the UCSC Genome Browser with the RNASeq tracks [64] to confirm the alternative splicing event. We then filtered these confirmed pairSum=9 events for those where there was a >20% average delta PSI in the 9 pairwise comparisons. Table 1 summarizes the number of confirmed alternative splicing events meeting these criteria in each strain comparison.

**Table 1.** Summary of Splicing Changes.

| pairSum = 9 with<br>Minimum ΔPSI = 0.15<br>& Average ΔPSI >0.20 (n=9) | KIN17 (K23N) | KIN17 (M107I) | PRCC (I371F) | PRCC (null) |
| --- | --- | --- | --- | --- |
|  | <u>SZ340 vs. SZ345</u> | <u>SZ340 vs. SZ355</u> | <u>SZ340 vs. SZ346</u> | <u>SZ340 vs. SZ356</u> |
| Alternative 5' Events | 4 | 3 | 69 | 90 |
| Alternative 3' Events | 108 | 24 | 1 | 35 |
| Skipped Exons | 7 | 2 | 0 | 5 |
| Retained Introns | 2 | 0 | 2 | 1 |
| Multi Skipped Exons | 0 | 0 | 0 | 0 |
| Mutually Exclusive Exons | 1 | 0 | 0 | 0 |
| Alternative First Exons | 5 | 1 | 0 | 5 |
| Alternative Last Exons | 7 | 1 | 0 | 1 |

### PRCC(I371F) and PRCC(null) promote usage of 5’ /UU splice sites and degenerate 5’ /GU splice sites throughout the *C. elegans* transcriptome

Using the stringent criteria described above, we were able to identify multiple examples of changes to 5’ splicing in the presence of PRCC mutations. In PRCC(I371F) and PRCC(null), we found, respectively, 34 and 46 examples of introns where mutant strains promote usage of a downstream /UU splice site over an adjacent /GU splice site (Fig 6B). This type of intron start of /G/UU 5’ splice site is similar to the *unc-73* splice site choice competition assays. Many of the introns affected by PRCC(I371F) are also affected by PRCC(null) (Fig 6E). These introns are enriched for an A in the 4th position of the intron immediately following the invariant GUU (Fig 6B).

**Figure 6.**
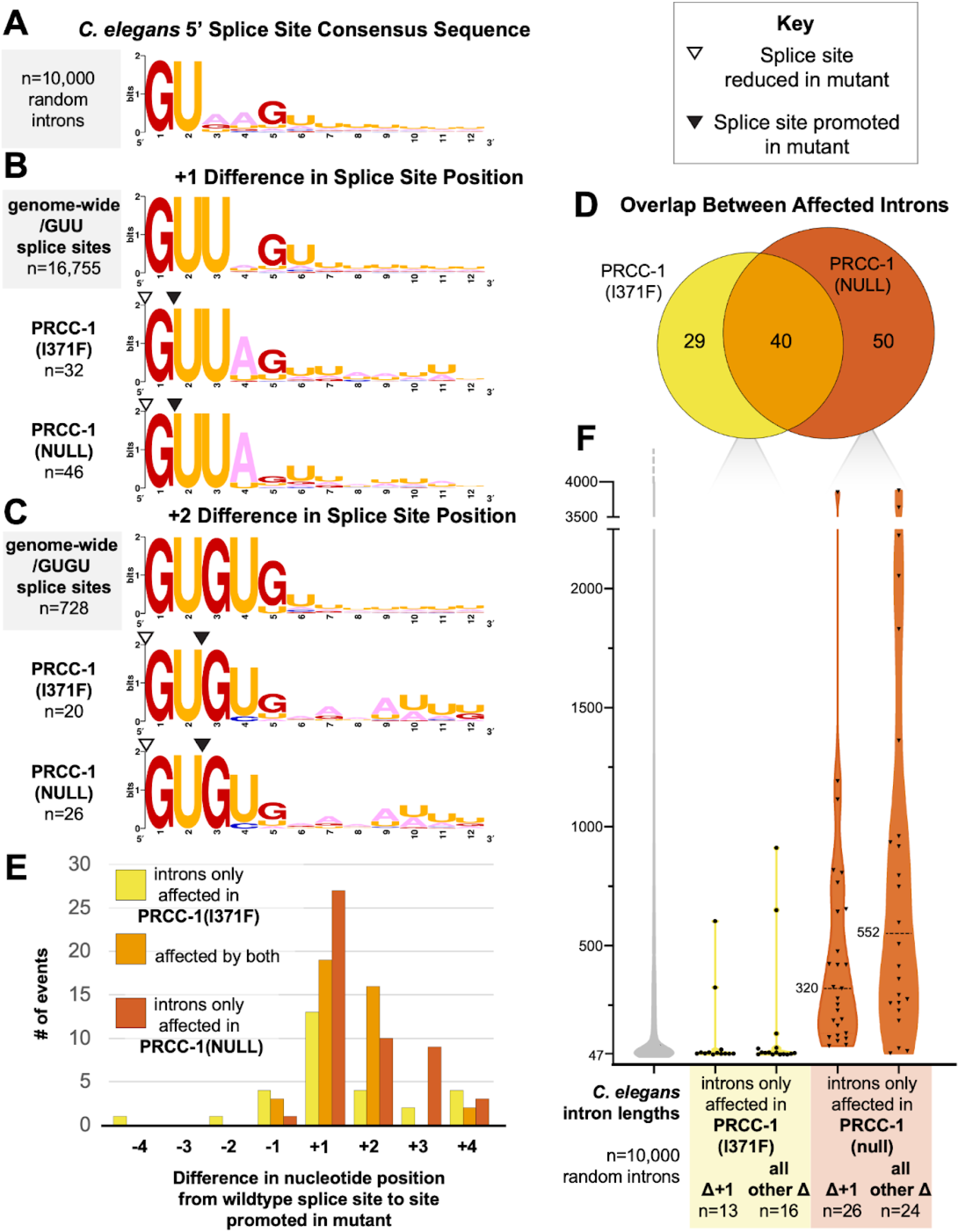
Throughout the genome, mutations in PRCC increase usage of /UU 5’ splice sites and /GU 5’ splice sites lacking other features. **(A)** Sequence logo showing the consensus sequence for the 5’ end of 10,000 randomly chosen *C.elegans* introns. **(B)** Sequence logo of introns that are differentially spliced in PRCC mutant backgrounds and follow the /G/UU splicing pattern seen in *unc-73(e936)* compared to all annotated introns that begin with /GUU. The splice site promoted in mutant is +1 nucleotides from the position of the predominant/GU splice site. Splice sites are indicated by triangles, as described in the key. **(C)** Sequence logo of introns that are differentially spliced in PRCC mutant backgrounds in which the splice site promoted in mutant is +2 nucleotides from the position of the predominant/GU splice site. Splice sites are indicated by triangles, as described in the key. **(D)** Euler diagram enumerating the overlap between affected introns differentially spliced in the presence of the two PRCC alleles. **(E)** Most splice sites promoted by the PRCC alleles are either one or two nucleotides downstream of the predominant splice site. Frequency and direction of nucleotide shift between the splice site favored in wild type, and the splice site promoted in PRCC mutant. **(F)** Violin plot showing the lengths of introns affected only in a given suppressors group. The five violins correspond to: 10,000 random wild-type C. *elegans* RefSeq introns, the subset of 13 introns in PRCC(I371F) in which the splice site promoted was at a /UU splice site 1 nucleotide downstream from the predominant splice site, the 16 affected introns in that same strain that did not follow +1 pattern, the subset of 26 introns in PRCC(null) in which the splice site promoted was at a /UU splice site 1 nucleotide downstream from the predominant splice site, the 24 affected introns in that same strain that did not follow +1 pattern. These groups of introns have median lengths of 47, 48, 51, 320 and 552 nucleotides, respectively.

In PRCC(I371F) and PRCC(null), background, we also found 37 and 44 instances, respectively, of events where the alternative 5’ splice site promoted in the presence of PRCC mutations were at /GU dinucleotides, either 2,3, or 4 nucleotides away from the wild-type /GU dinucleotide. Most of these shifted downstream (Fig 6E). A substantial portion of the introns affected by the PRCC-1(null) were also affected by the point mutation in PRCC(I371F) (Fig 6D). Surprisingly, despite the similarity between the splicing phenotypes observed in our *unc-73(e936)*-based splice site competition assays for both PRCC and KIN17 mutations, we found negligible examples of changes to 5’ splice site choice at endogenous introns in the presence of either of the two KIN17 mutant alleles.

### PRCC-1 null 5’ affects longer introns, especially in the case of non-GUU introns

We were interested in the group of introns affected by PRCC mutations, so we looked at the lengths of introns, and flanking exons. Despite the overlap between affected introns, the average intron length for each group is very different. Because rare, very long introns can exert a strong influence on averages, we report the median intron length. In order to focus more on the relative contribution to median intron length in each category, we removed events in common and looked at the lengths of introns unique to each dataset (Fig 6D). While the median intron length for /UU and /GU alternative splice sites promoted in PRCC(I371F) background is similar to the overall median intron length in *C. elegans* of 51 nucleotides [25], the median intron length of PRCC(null) promoted alternative introns for both /UU and /GU introns is much longer, with a median length of 320 and 552 nucleotides respectively (Fig 6F).

### KIN17(K23N), KIN17(M107I) and PRCC(null) promote usage of weak upstream 3’ splice sites throughout *C. elegans* transcriptome

Even more surprising than the inability of KIN17 mutations to change global 5’ splice site choice, is the ability of those same mutations, identified in a screen for modifiers of 5’ splice choice, to affect 3’ splice site choice. The 3’ splice sites promoted in the presence of these KIN17 mutations were highly degenerate sites (Fig 7A), mostly located 6 or 9 base pairs away and unidirectionally upstream of the adjacent consensus 3’ splice sites (Fig 7B). We found 108 examples of alternative 3’ss usage in KIN17(K23N), 24 examples in KIN17(M107I) and 35 examples in the PRCC(null) (Table 1). We only found one example of 3’ changes in PRCC(I371F). Most of the intron events identified in KIN17(M107I) were also represented in the KIN17(K23N) events (Fig 7C). We found only 5 unique examples of PRCC(null) mutations affecting 3’ splice site choice that are not shared with the KIN17 mutant strains.The unidirectional shift to a poor consensus upstream 3’ss is similar to developmentally regulated alternative splicing events in which cells in the *C. elegans* germline show more splicing to an upstream, poor consensus alternative 3’ss relative to somatic cells (Ragle et al., 2015). In that study, 203 alternative 3’SS events were identified as being developmentally regulated; 49 of those alternative 3’ splicing events overlap with the alternative 3’ splicing events identified in PRCC and KIN17 mutants (Fig 7D).

**Figure 7.**
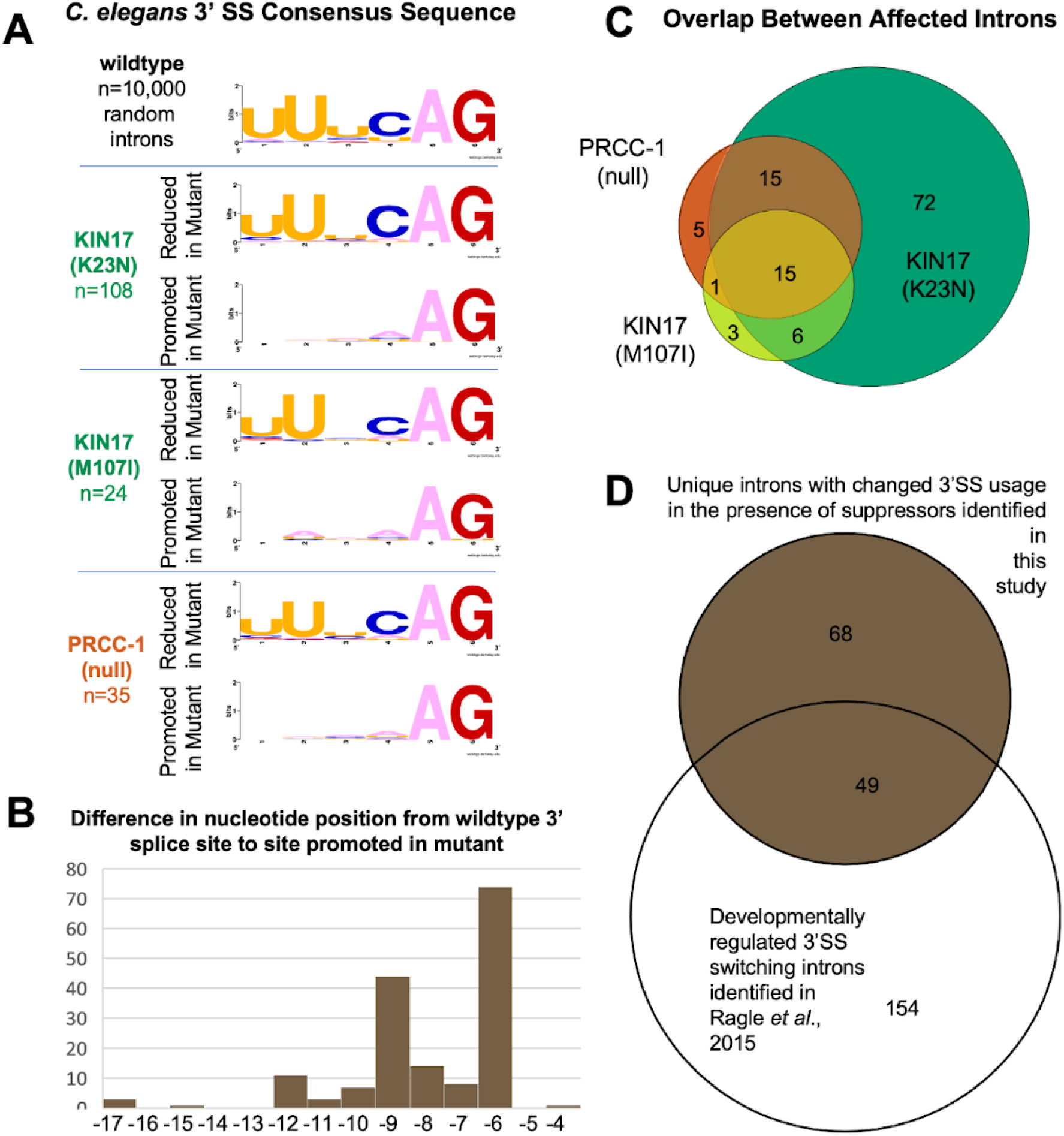
Mutations in KIN17 and PRCC(null) promote usage of 3’ splice sites with minimal consensus sequence, upstream of 3’ canonical splice sites. **(A)** *C. elegans* 3’ splice site consensus sequence for 10,000 random wild-type introns, followed by the consensus sequence of the splice sites that were reduced in the mutant strains and then the consensus sequence of the splice sites that were promoted in the strains with mutations in KIN17(K23N), KIN17(M107I) and PRCC(null) respectively. **(B)** Most splice sites whose usage increases in the presence of KIN17(K23N), KIN17(M107I) and PRCC(null) are either 6 or 9 nucleotides upstream of the predominant wild-type splice site. Frequency of nucleotide shift between the splice site favored in wild type, and the splice site promoted in PRCC mutant. **(C)** Euler diagram shows extent of overlap between intronic events with changed 3’ splice site choice in KIN17(K23N), KIN17(M107I), and PRCC(null). **(D)** Euler diagram shows extent of overlap between all unique intronic events with changed 3’ splice site choice in this study, compared to the developmentally regulated 3’SS switching previously identified by our lab, in which certain introns show a shift towards usage of an alternative upstream 3’ SS in the germline, which has minimal consensus sequence aside from an AG dinucleotide at the end of the intron [25].

## Discussion

This work represents the first direct demonstration that KIN17 and PRCC have a role in splice site choice. Prior to this manuscript, KIN17 was classified in the Spliceosome Database under “misc. proteins found irregularly with spliceosomes” (http://spliceosomedb.ucsc.edu/proteins/11606, accessed 3/22/2021), and had been primarily studied for roles in DNA damage repair and cancer, not splicing. We report here that mutations in the N-terminal unstructured region (K23N) and in the winged helix (M107I) of KIN17 promote usage of an unusual /UU 5’ splice site mostly downstream of an adjacent /GU splice site (Figs 1 and 6), and, even more surprisingly, those same mutations change 3’ splice site choice at over a hundred native loci, promoting degenerate splice sites upstream of canonical 3’splice sites (Fig 7). Excitingly, while we were preparing this manuscript, a structure of the pre-B^act^2 spliceosome was published [15], with KIN17 modeled in this transient intermediate near what will become the active site later on in the splicing cycle (Fig 3). The loop and the 3_10_ turn of the winged helix are positioned facing the active site, though the M107 residue points into the globular core of the winged helix, not outward. This leads us to hypothesize that the M107I mutation repositions nearby outward facing residues such as the highly conserved nearby aromatic residues: histidine 104, histidine 106, and tryptophan 112. Townsend *et al.*, hypothesize an early transient role in spliceosome assembly for KIN17, proposing that KIN17 prevents components of the spliceosome, including PRP-8 and BBR2, from prematurely entering the B^act^ conformation. In light of our result showing significant alterations to 3’ splice sites, we hypothesize that KIN17 either has an additional later role in the splicing cycle, or that the premature assembly of B^act^ leads to later acceptance of an upstream branch point corresponding to a degenerate 3’ splice site, as the branch point itself is not yet positioned for catalysis in B^act^. This demonstration of KIN17 as a bona fide splicing factor may potentially point to a closer association between pre-mRNA splicing and DNA damage repair than is currently understood. PRP19 is a multifunctional ubiquitin ligase known to be a component of both spliceosomal and DNA damage repair complexes [65], and a recent study showed that U1snRNP and components of the DNA damage response compete for binding at human 5’ splice sites [66]. As both splicing and DNA damage repair require the recognition, cutting and joining of nucleic acid chains, it may not be too surprising that they share some factors in common.

Prior to our studies, PRCC had a firmer association to the spliceosome, identified as a factor in B^act^ complexes through Yeast two-hybrid and mass spectrometry experiments [13,60], but no functional role had been identified nor had it been modeled into any metazoan spliceosomal structures (there is no *S. cerevisiae* homolog of this factor). We report that a I371F point mutation, located in the 9-residue-long region in the C-terminus of PRCC that is identical between worms and humans, changes 5’ splice site choice at native loci, and PRCC(null) promotes both noncanonical downstream 5’ splice sites and noncanonical upstream 3’ splice sites. It is possible that PRCC is serving a different function in *C. elegans* than it does in other organisms; the “proline rich-region” of PRCC most often found in oncogenic fusions is noticeably proline-poor in the *C. elegans* homolog relative to humans. The identification of a suppressor point mutation in a conserved region of the C-terminus points to a potential key region for splicing control.

The discovery of this new class of suppressors of *unc-73(e936)* cryptic splicing has led us to think about the splice site like a piece of evidence in a criminal case, held by “escorts” which shuttle the precise genetic landmarks through dramatic conformational changes. Each escort of the 5’ splice site, must by nature, hold it reversibly. Therefore, slipping or disengagement are possible while the 5’ss is in the custody of a snRNP or protein factor guardian, especially when the pre-mRNA is under tension from helicases or other components of the spliceosome. If we follow the chain of custody, we expect that translocations and changes of possession, are likely to be inflection points where alterations to splice site identity, relative to the initial identification by early factors, are more likely. Some factors capable of affecting splice site choice may assist during those vulnerable moments in the splicing cycle. When an escort repositions or lets go entirely, these factors may make nucleotide shifts less likely. We see in the presence of the suppressors identified in this study, that the spliceosomal components are choosing degenerate splice sites. The positions we have identified in KIN17 and PRCC may serve to prevent such slips in wild type during vulnerable points in the chain of custody.

These mutations display a different splicing phenotype from previously identified suppressors. Instead of the predictable reduction of the distal +23 site and relatively even increase in usage of both splice sites of the doublet observed in factors previously identified (Fig 1D) ([16,17]), this new class of type III suppressors displays a sharp change in the ratio of usage of the two adjacent splice sites of the doublet of adjacent splice sites, with the downstream /UU site promoted over the adjacent /GU site (Fig 1D). This effect is seen with or without other nearby cryptic /GU splice sites (Fig 1 and Fig 5B), and can be replicated at a downstream location (Fig 5D). We believe this difference between Class III suppressors and previously identified suppressors supports the idea that these factors act at a different point in the splicing cycle. The first U1 dependent step of 5’ss identification can be thought of like the coarse focus on a microscope, and the class II suppressors can be thought of as mutations to factors that maintain the general region of the identified splicing target. In later steps after U1 has left, we can think of the maintenance of the 5’ss as a more “fine focus” function, perhaps related to U6 identification of the 5’ss [67] and the class III suppressors are mutations that alter the ability of the spliceosome to maintain the fine focus of the splice site that will be used in chemistry, an effect that is consistent with the duplicated doublet switching result (Fig 5D).

We were surprised that this genetic screen for factors that affect 5’ splice site choice identified factors capable of affecting both 5’ and 3’ splice site choice. We were further surprised that despite the similar splicing phenotypes displayed in *unc-73*-based reporters of splice site choice, there were differences in how suppressors affected global splicing in an NMD background. Both PRCC suppressors affected global 5’ss choice, promoting usage of non-consensus 5’ss downstream of canonical 5’ss, especially at long introns, but neither of the two KIN17 suppressors affected global 5’ splicing. Both KIN17 suppressors affected global 3’ss, promoting usage of non consensus 3’ss upstream of canonical 3’ss, as did PRCC(null), but not PRCC(I371F). No suppressors identified in this screen promoted significant numbers of other splicing alterations, such as alternative first or last exons, exon or intron inclusion or skipping (Table 1). All effects observed were local, usually shifting 5’ splice site choice by 1 or 2 nucleotides downstream, and 3’ss choice usually by either 6 or 9 nucleotides upstream.

This preference for upstream non consensus 3’ss reminded us of the tissue-specific 3’ss switching identified by our lab [25]. In addition, many of the upstream AGs found to be prefered in germline tissue relative to somatic tissue are also preferred in KIN17 and PRCC mutants relative to wild type (Fig 7D). Mutations in various parts of the spliceosome act on some of these same splice sites (this work and unpublished data). A simple interpretation of this overlap is that there are a relatively small number of ambiguous adjacent 3′ splice sites present in the genome, and that there are multiple mechanisms involved in making the distinction. Our work shows that the genetic probing of subtle changes to splice site choice by which we have studied various 5′ mechanisms can also be used to study 3′ splice site choice mechanisms. Despite these mutations affecting the choice made during the second splicing reaction, we should not take these results as strong evidence that KIN17 and PRCC still are present and functioning in late spliceosomal complexes. KIN17 and PRCC may be influencing the choice before the second step occurs. One possibility is that these mutations alter branchpoint choice, and then the alternate 3′ splice site choice is a secondary effect. Another possibility is that these mutations alter the form of the spliceosome to affect 3’ splice site choice in a way that persists during the second step, even after the proteins themselves are removed.

## Methods

Full step-by-step protocols of many of the methods described below have been deposited at https://dx.doi.org/10.17504/protocols.io.p9kdr4w.

### Growth Conditions

*C. elegans* were maintained at 20°C on nematode growth medium (NGM) agar plates inoculated with OP50 *E. coli.* Strains were discovered in the suppressor screen, genetically engineered using CRISPR mutagenesis, created by doing genetic crosses, or obtained from the *C. elegans* Gene Knockout Consortium [61].

### *C. elegans*strains

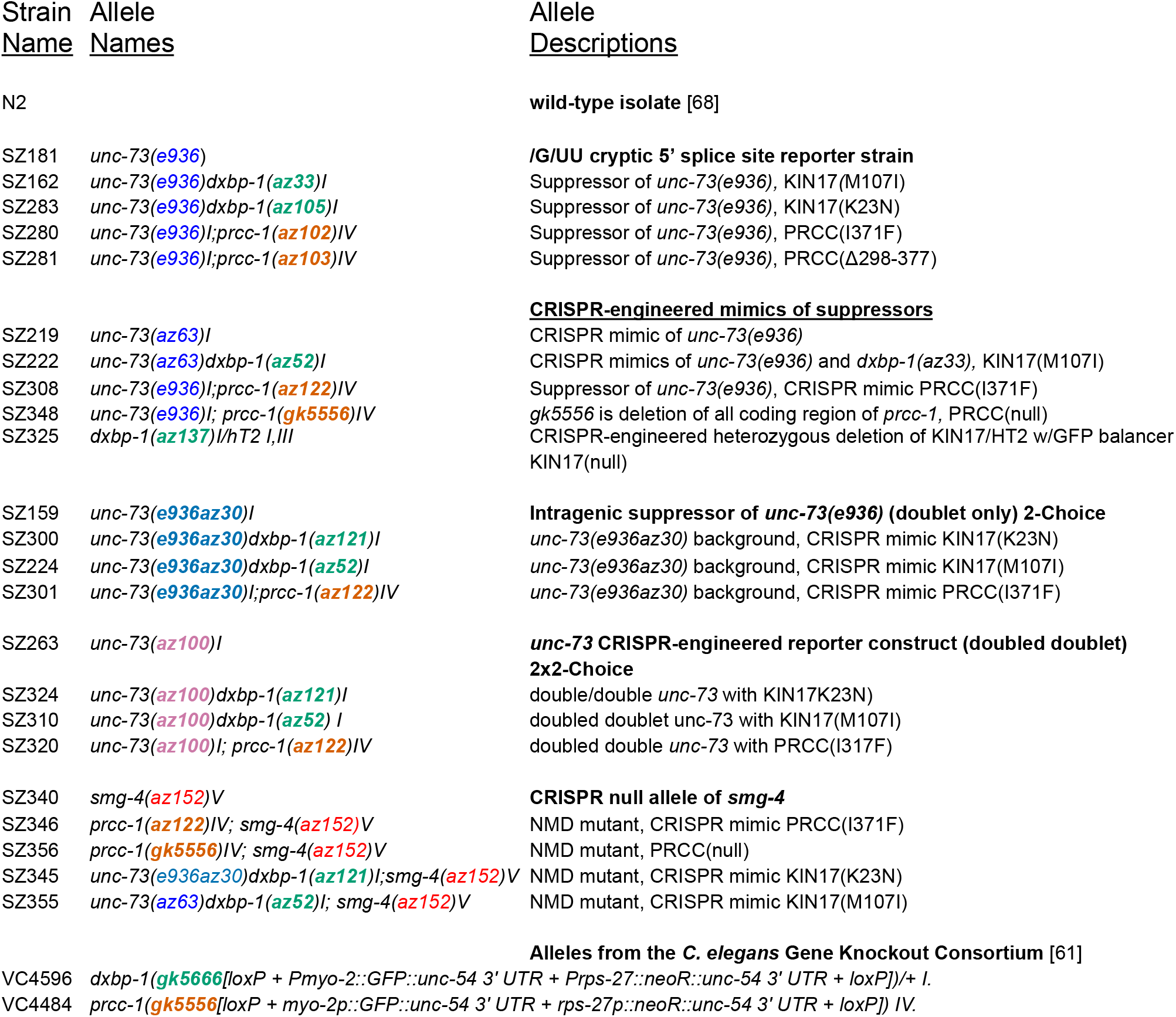

### Mutagenesis and identification of putative suppressed strains

Age-synchronized uncoordinated *unc-73(e936)* hermaphrodites in gametogenesis, larval stage L4, were soaked in 0.5mM N-nitroso-N-ethylurea (ENU) as previously described [16]. After extensive washing, four animals were placed at the edge of an OP50 *E. coli*-seeded 10cm NGM-agar plate, for 500 plates, and allowed to self-propagate. NGM plates were maintained at 20°. Whereas the *unc-73(e936)* animals’ movement defects confine them in place, after 8 days, suppressed F2 animals are able to crawl away from the crowded pile of uncoordinated animals, and are identifiable by their improved locomotion on the far side of the plate.

### Identification of extragenic splicing suppressors

The *unc-73* gene in suppressed lines from this screen was sequenced to distinguish between extragenic and intragenic suppressors; one of these intragenic suppressors, *unc-73(e936az30)* is used in this study (Fig 5A). Remaining extragenic suppressor alleles were mapped to chromosomes using a strategy described in [26,69]. Briefly, each suppressor strain identified in the genetic screen was crossed against a polymorphic Hawaiian isolate CB4856 and uncoordinated F2 animals that continued to have only uncoordinated offspring were recovered. These new Unc strains were then screened for regions that are homozygous for snip-SNP markers as described by [26]. Approximately 20 uncoordinated strains for each extragenic suppressor strain outcrossed to the Hawaiian strain were recovered and DNA extracted and combined. For each chromosomal region, we expected to see a mix of Hawaiian and Bristol N2 single nucleotide polymorphisms (SNPs), except in the region linked to the suppressor mutation, where we expect to see 100% Hawaiian SNPs (loss of the suppressor in the N2 background) and in the region of *unc-73* where we expect to see 100% N2 SNPs (the uncoordination allele is in the N2 background). Using this approach, we were able to narrow down the suppressors to approximately one third of the length of a chromosome. At the same time, the suppressor strains were backcrossed two times to the N2 wild-type strain, and then we performed high-throughput genomic sequencing of the suppressor strains. We used STAR [70] to map those sequences back to the *C. elegans* genome. Diploid SNPs relative to the original N2 strain were identified using GATK [71]. The snpEff tool [72] was used to identify SNPs within genes in the chromosomal region identified by the Hawaiian strain mapping. That list of putative suppressors was cross-referenced to the Jurica lab Spliceosome database, [73], (http://spliceosomedb.ucsc.edu/) and candidate spliceosome-associated genes and RNA binding proteins in the delimited genomic region were chosen for further analysis. The suppressor allele identity was verified by *de novo* re-creation of each putative suppressor allele using CRISPR/Cas9 genome editing, and those resulting in both suppression of the movement defect and molecular changes in splicing were identified as *bona fide* suppressors.

### CRISPR/Cas9 Genome editing

Cas9 guides were chosen from the CRISPR guide track on the UCSC Genome Browser *C. elegans* reference assembly [64,74,75] and crRNAs were synthesized by Integrated DNA Technologies (www.idtdna.com). Cas9 CRISPR RNA guides were assembled with a standard tracrRNA; these RNAs were heated to 95°C and incubated at room temperature to allow joining. The full guides were then incubated with Cas9 protein to allow for assembly of the CRISPR RNA complex [76]. That mix, along with a single stranded repair guide oligonucleotide was then micro-injected into the syncytial gonad of young adult hermaphrodite animals. A *dpy-10(cn64)* co-CRISPR strategy was used to identify F1 animals showing homologous recombination CRISPR repair in their genomes [77]. Silent restriction sites were incorporated into repair design so that mutations could be easily tracked by restriction digestion of PCR products from DNA extracted from single worms. Injected animals were moved to plates in the recovery buffer [76], allowed to recover for 4 hours, and moving worms were plated individually. F1 offspring were screened for the *dpy-10(cn64)* dominant roller (Rol) co-injection marker phenotype. F1 Rol animals were plated individually, allowed to lay eggs, and then the adult was removed and checked for allele of interest by PCR followed by restriction enzyme digestion and gel electrophoresis. If an F1 worm showed the presence of a heterozygous DNA fragment matching the programmed restriction site, non-rollers in the F2 generation of that worm were screened by electrophoresis of digested PCR products. Individuals that had lost the co-injection marker, but were homozygous for the allele of interest were retained and sequenced at the gene of interest to verify error-free insertion of sequences guided by the repair oligo.

### CRISPR Sequences

Suppressor mutations are bold and capitalized

Silent mutations for preventing recut or for restriction sites are capitalized or starred

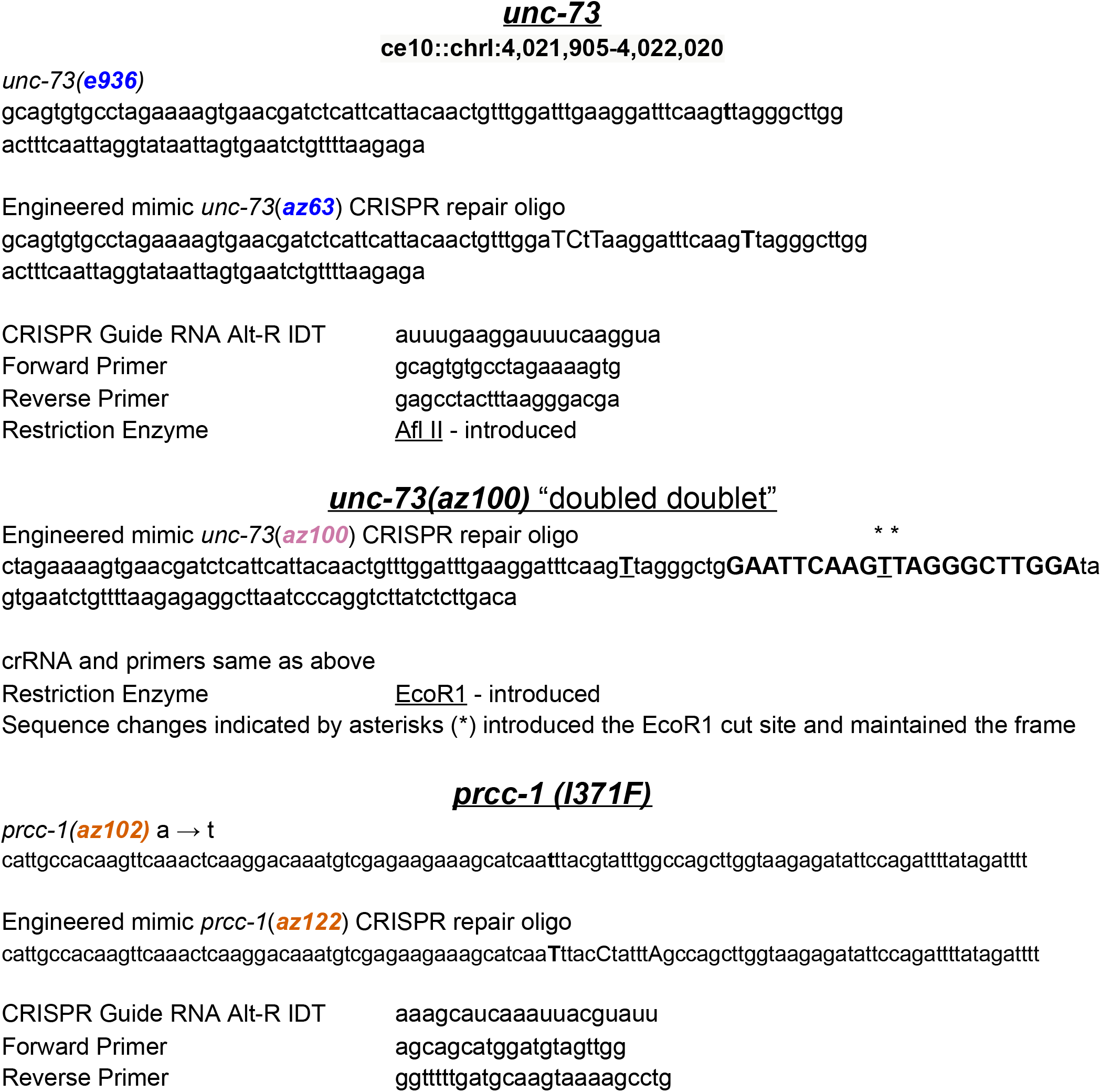
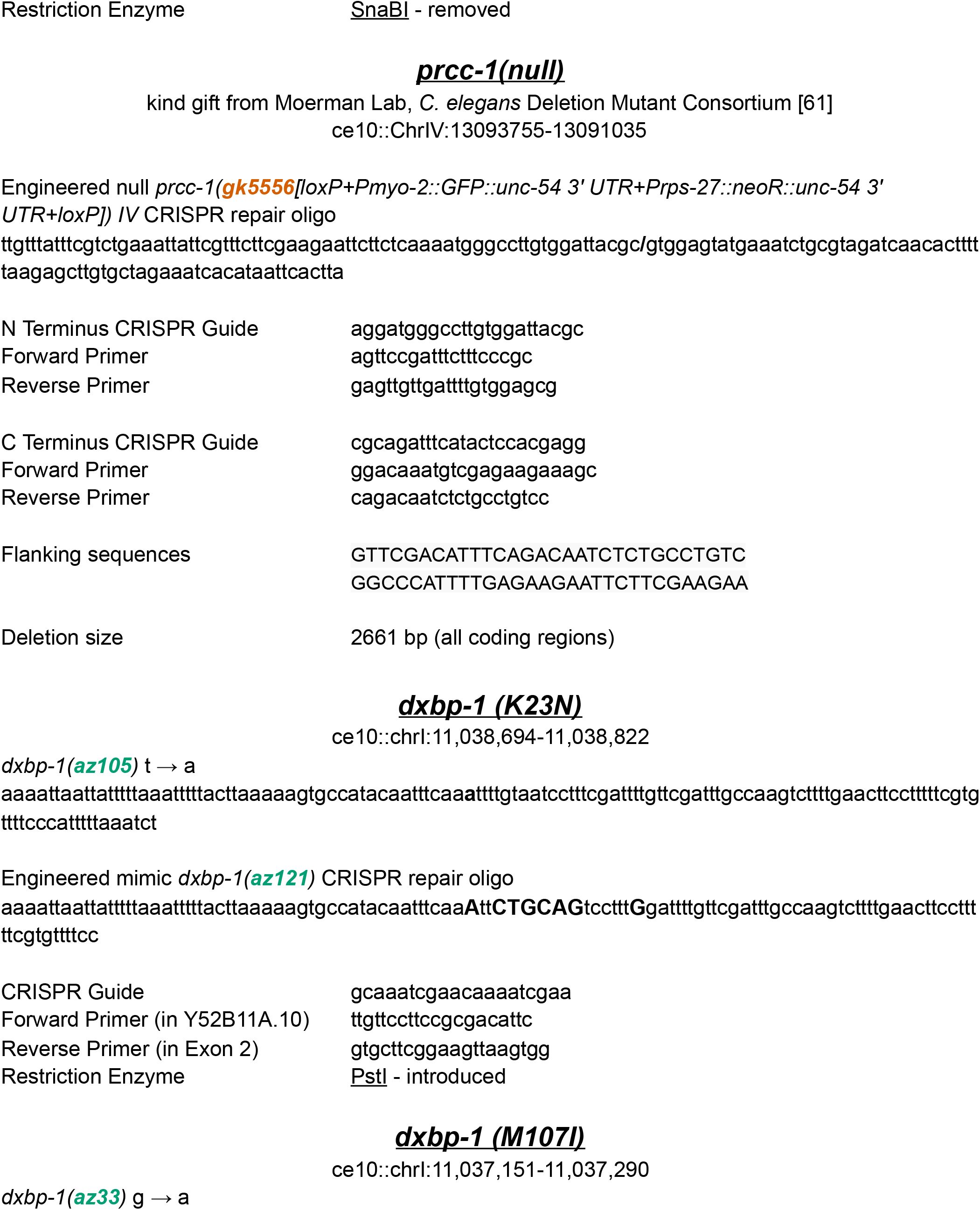
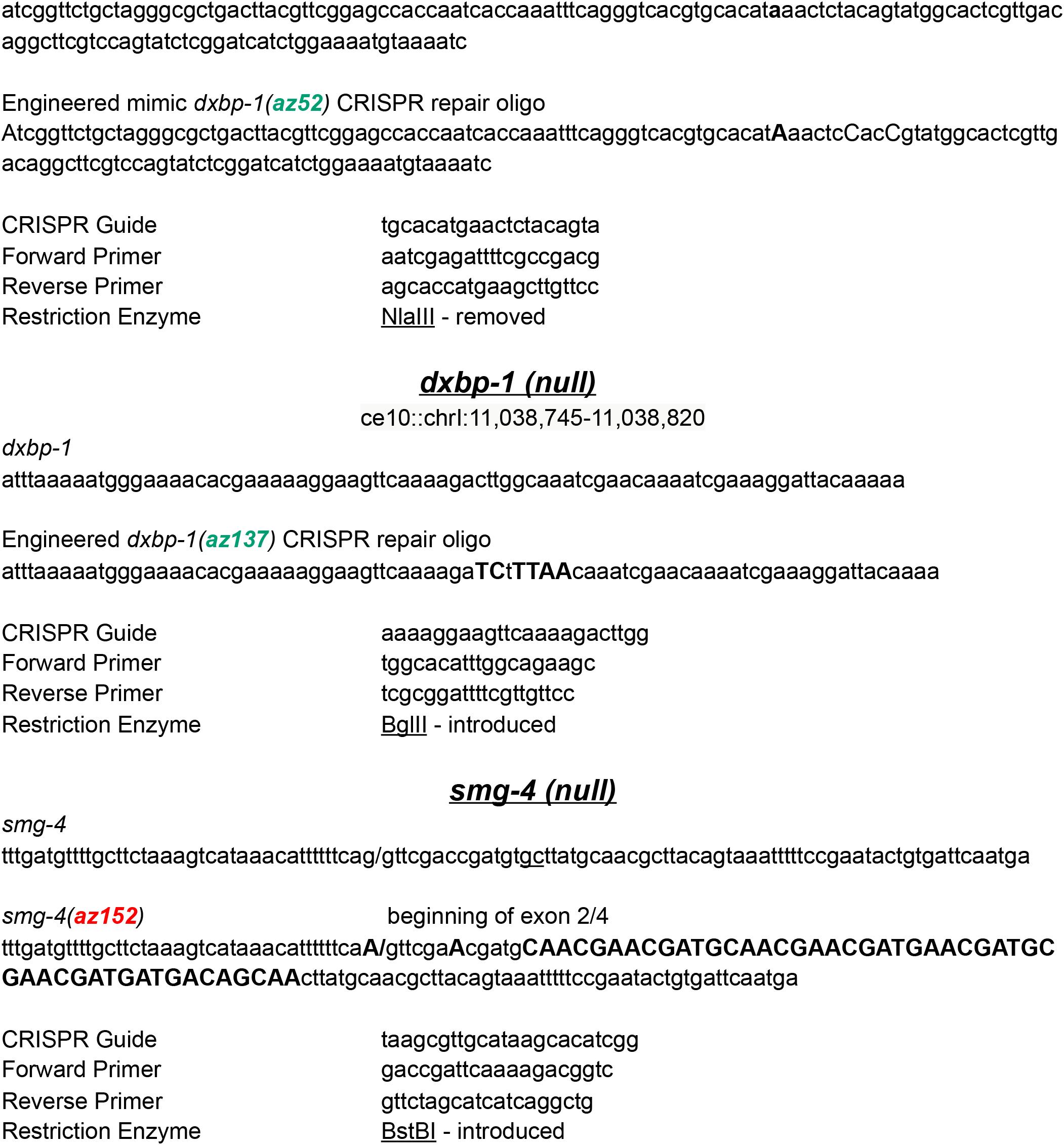

### RNA extraction, cDNA production and PCR amplification

RNA from indicated strains was extracted from mixed stage populations of animals using TRIzol reagent (Invitrogen), then alcohol precipitated. Total RNA was reverse transcribed with gene-specific primers using SuperScript III (ThermoFisher) or AMV reverse transcriptase (Promega). cDNA was PCR-amplified for 25 cycles with 5’-Cy3-labelled reverse primers (IDT) and unlabeled forward primers using either Taq polymerase or Phusion high-fidelity polymerase (NEB). PCR products were separated on 40cm tall 6% polyacrylamide denaturing gels and then visualized using a Molecular Dynamics Typhoon Scanner. Band intensity quantitation was performed using ImageJ software (https://imagej.nih.gov/ij/).

### RNASeq

Triplicate total RNA isolations were done for each strain, and mRNA sequencing libraries were prepared for each RNA isolation by RealSeq Biosciences (Santa Cruz, CA). 75 x 75 paired- end reads were obtained on a Novaseq 6000 sequencer, with 9 libraries combined in a lane. RNA-seq results were trimmed, subjected to quality control, and two-pass aligned to UCSC Genome Browser *C. elegans* reference assembly (this earlier assembly release was used to facilitate comparison to previous RNA-seq datasets obtained by our lab) using a modified version of STAR [70]. The standard version of STAR, in addition to the canonical GU/AG intron motif, supports GC/AG and AU/AC motifs for the 5’ and 3’ splice sites. Because *C. elegans* does not have minor spliceosomes with AU at the 5’ end of introns, we modified the STAR source code to use UU/AG as the third motif in place of AU/AC. Furthermore, we ran STAR with parameters that adjusted the default “scoreGapATAC” (effectively scoreGapUUAG in our modified version of STAR) junction penalty from −8 to 0 so that the program would treat UU/AG spliced introns with the same scoring as GU/AG introns.

### High Stringency ΔPSI Analysis

Alternative 5’ (A5) and alternative 3’ (A3) splicing events found in the STAR mappings of all of the libraries were identified and filtered for those introns with at least 5 reads of support (total across all samples) and a maximum of 50 nucleotides between the alternative ends (either 5’ or 3’ respectively). In addition, alternative first exon (AF), alternative last exon (AL), skipped exon (SE), retained intron (RI), mutually exclusive exon (MX) and multiple skipped exon (MS) events were derived from the Ensembl gene predictions Archive 65 of WS220/ce10 (EnsArch65) using junctionCounts “infer pairwise events” function (https://github.com/ajw2329/junctionCounts). The percent spliced in (PSI) in each sample was derived for all of these events using junctionCounts. Pairwise differences in PSI between samples for the above events were calculated. Alternative splicing events with a minimum 15% ΔPSI were included for further consideration. Each strain had 3 biological replicates, therefore between any two strains, a total of nine pairwise comparisons were possible between each suppressor strain and the SZ340 *smg-4* comparison strain for each alternative splicing event. For each suppressor strain, only alternative splicing events that showed a change in the same direction >15% ΔPSI compared to the smg-4 control in all nine pairwise comparisons (pairSum=9) were considered. Those events with a mean ΔPSI >20% across the 9 comparisons were included for further consideration. The reads supporting that alternative splice site choice event were then examined by eye on the UCSC Genome Browser *C. elegans* reference assembly to ensure that the algorithmically flagged events looked like real examples of alternative splice site choice. Supplemental table 1 has the chromosomal location, PSI measurements and notes for all alternative splicing events that fit these criteria.

### Sequencing Data Access

Raw mRNA sequencing data for 15 libraries in fastq format, along with .gtf files for all analyzed alternative splicing events, are available in fastq format at the NCBI Gene Expression Omnibus (GEO - https://www.ncbi.nlm.nih.gov/geo/) accession GSE178335.

### Consensus Motifs

Consensus motifs were created using WebLogo [78]; https://weblogo.berkeley.edu/logo.cgi.

### Multiple Sequence Alignments

Multiple sequence alignments were generated using the EMBL-EBI Clustal Omega MSA webtool [79]; https://www.ebi.ac.uk/Tools/msa/clustalo/).

## Acknowledgments

We thank Noel Ng and Eimy Castellanos for technical assistance. We are grateful to Michael Doody, Melissa Jurica, Manny Ares, Harry Noller and Jordan Eizenga for helpful discussions. We thank Josh Arribere for assistance identifying mutant alleles from sequencing data and Guillaume Chanfreau for the suggestion of using Cy3-labeled PCR primers.

Research in the Zahler lab was supported by a grant from the National Science Foundation (MCB-1613867 to AMZ) and currently by a grant from the National Institutes of Health (5R01GM135221 to AMZ). JMNGL was supported by the UCSC MCD Graduate Training Grant (5T32GM133391).

## Author Contributions

Jessie M.N.G. Lopez 1

Conceptualization
Formal Analysis
Investigation
Methodology
Project Administration
Validation
Visualization
Writing - Original Draft Preparation
Writing - Review & Editing

Kenneth Osterhoudt 1

Investigation
Methodology
Writing - Review & Editing

Catiana Holland Cartwright 1

Formal Analysis
Investigation
Validation
Visualization

Sol Katzman 2

Data Curation
Investigation
Software
Writing - Review & Editing

Alan M. Zahler 1

Conceptualization
Funding Acquisition
Investigation
Methodology
Project Administration
Resources
Supervision
Writing - Review & Editing
Validation

## Notes

https://www.ncbi.nlm.nih.gov/geo/query/acc.cgi?acc=GSE178335

### Competing Interest Statement

The authors have declared no competing interest.

## References

1. Staley JP, Guthrie C. Mechanical devices of the spliceosome: motors, clocks, springs, and things. Cell. 1998;92: 315–326.

2. Wilkinson ME, Charenton C, Nagai K. RNA Splicing by the Spliceosome. Annu Rev Biochem. 2020;89: 359–388.

3. Herzel L, Straube K, Neugebauer KM. Long-read sequencing of nascent RNA reveals coupling among RNA processing events. Genome Res. 2018;28: 1008–1019.

4. Rinke J, Appel B, Blöcker H, Frank R. The 5′-terminal sequence of U1 RNA complementary to the consensus 5′ splice site of hnRNA is single-stranded in intact U1 snRNP particles. Nucleic acids. 1984. Available: https://academic.oup.com/nar/article-abstract/12/10/4111/1137838

5. Wong MS, Kinney JB, Krainer AR. Quantitative Activity Profile and Context Dependence of All Human 5’ Splice Sites. Mol Cell. 2018;71: 1012–1026.e3.

6. Zorio DA, Blumenthal T. Both subunits of U2AF recognize the 3’ splice site in Caenorhabditis elegans. Nature. 1999;402: 835–838.

7. Berglund JA, Abovich N, Rosbash M. A cooperative interaction between U2AF65 and mBBP/SF1 facilitates branchpoint region recognition. Genes Dev. 1998;12: 858–867.

8. Stenson PD, Mort M, Ball EV, Evans K, Hayden M, Heywood S, et al. The Human Gene Mutation Database: towards a comprehensive repository of inherited mutation data for medical research, genetic diagnosis and next-generation sequencing studies. Hum Genet. 2017;136: 665–677.

9. Sterne-Weiler T, Howard J, Mort M, Cooper DN, Sanford JR. Loss of exon identity is a common mechanism of human inherited disease. Genome Res. 2011;21: 1563–1571.

10. Glidden DT, Buerer JL, Saueressig CF, Fairbrother WG. Hotspot exons are common targets of splicing perturbations. Nat Commun. 2021;12: 2756.

11. Dyle MC, Kolakada D, Cortazar MA, Jagannathan S. How to get away with nonsense: Mechanisms and consequences of escape from nonsense-mediated RNA decay. Wiley Interdiscip Rev RNA. 2020;11: e1560.

12. Malca H, Shomron N, Ast G. The U1 snRNP base pairs with the 5’ splice site within a penta-snRNP complex. Mol Cell Biol. 2003;23: 3442–3455.

13. Agafonov DE, Kastner B, Dybkov O, Hofele RV, Liu W-T, Urlaub H, et al. Molecular architecture of the human U4/U6.U5 tri-snRNP. Science. 2016;351: 1416–1420.

14. Maroney PA, Romfo CM, Nilsen TW. Functional recognition of 5’ splice site by U4/U6.U5 tri-snRNP defines a novel ATP-dependent step in early spliceosome assembly. Mol Cell. 2000;6: 317–328.

15. Townsend C, Leelaram MN, Agafonov DE, Dybkov O, Will CL, Bertram K, et al. Mechanism of protein-guided folding of the active site U2/U6 RNA during spliceosome activation. Science. 2020. p. eabc3753. doi:10.1126/science.abc3753

16. Dassah M, Patzek S, Hunt VM, Medina PE, Zahler AM. A genetic screen for suppressors of a mutated 5’ splice site identifies factors associated with later steps of spliceosome assembly. Genetics. 2009;182: 725–734.

17. Mayerle M, Yitiz S, Soulette C, Rogel LE, Ramirez A, Ragle JM, et al. Prp8 impacts cryptic but not alternative splicing frequency. Proc Natl Acad Sci U S A. 2019;116: 2193–2199.

18. Steven R, Kubiseski TJ, Zheng H, Kulkarni S, Mancillas J, Ruiz Morales A, et al. UNC-73 activates the Rac GTPase and is required for cell and growth cone migrations in C. elegans. Cell. 1998;92: 785–795.

19. Roller AB, Hoffman DC, Zahler AM. The allele-specific suppressor sup-39 alters use of cryptic splice sites in Caenorhabditis elegans. Genetics. 2000;154: 1169–1179.

20. Zahler AM, Tuttle JD, Chisholm AD. Genetic suppression of intronic+ 1G mutations by compensatory U1 snRNA changes in Caenorhabditis elegans. Genetics. 2004. Available: https://www.genetics.org/content/167/4/1689.short

21. Zahler AM, Rogel LE, Glover ML, Yitiz S, Ragle JM, Katzman S. SNRP-27, the C. elegans homolog of the tri-snRNP 27K protein, has a role in 5’ splice site positioning in the spliceosome. RNA. 2018;24: 1314–1325.

22. Patzek S, Hunt VM, Medina PE, Zahler AM. A genetic screen for suppressors of a mutated 5′ splice site identifies factors associated with later steps of spliceosome assembly. 2009. Available: https://www.genetics.org/content/182/3/725.short

23. Sievers F, Wilm A, Dineen D, Gibson TJ, Karplus K, Li W, et al. Fast, scalable generation of high-quality protein multiple sequence alignments using Clustal Omega. Mol Syst Biol. 2011;7: 539.

24. Carlier L, Couprie J, le Maire A, Guilhaudis L, Milazzo-Segalas I, Courçon M, et al. Solution structure of the region 51-160 of human KIN17 reveals an atypical winged helix domain. Protein Sci. 2007;16: 2750–2755.

25. Ragle JM, Katzman S, Akers TF, Barberan-Soler S, Zahler AM. Coordinated tissue-specific regulation of adjacent alternative 3’ splice sites in C. elegans. Genome Res. 2015;25: 982–994.

26. Davis MW, Hammarlund M, Harrach T, Hullett P, Olsen S, Jorgensen EM. Rapid single nucleotide polymorphism mapping in C. elegans. BMC Genomics. 2005;6: 118.

27. le Maire A, Schiltz M, Stura EA, Pinon-Lataillade G, Couprie J, Moutiez M, et al. A tandem of SH3-like domains participates in RNA binding in KIN17, a human protein activated in response to genotoxics. J Mol Biol. 2006;364: 764–776.

28. Despras E, Miccoli L, Créminon C, Rouillard D, Angulo JF, Biard DSF. Depletion of KIN17, a human DNA replication protein, increases the radiosensitivity of RKO cells. Radiat Res. 2003;159: 748–758.

29. Kannouche P, Pinon-Lataillade G, Tissier A, Chevalier-Lagente O, Sarasin A, Mezzina M, et al. The nuclear concentration of kin17, a mouse protein that binds to curved DNA, increases during cell proliferation and after UV irradiation. Carcinogenesis. 1998;19: 781–789.

30. Angulo JF, Mauffirey P, Pinon-Lataillade G, Miccoli L, Biard DSF. Putative Roles of kin17, a Mammalian Protein Binding Curved DNA, in Transcription. DNA Conformation and Transcription. pp. 75–89. doi:10.1007/0-387-29148-2_6

31. Mazin A, Timchenko T, Ménissier-de Murcia J, Schreiber V, Angulo JF, Gilbert de M, et al. Kin17, a mouse nuclear zinc finger protein that binds preferentially to curved DNA. Nucleic Acids Res. 1994;22: 4335–4341.

32. Kannouche P, Angulo JF. Overexpression of kin17 protein disrupts nuclear morphology and inhibits the growth of mammalian cells. J Cell Sci. 1999;112 (Pt 19): 3215–3224.

33. Biard DSF, Saintigny Y, Maratrat M, Paris F, Martin M, Angulo JF. Enhanced expression of the Kin17 protein immediately after low doses of ionizing radiation. Radiat Res. 1997;147: 442–450.

34. Masson C, Menaa F, Pinon-Lataillade G, Frobert Y, Radicella JP, Angulo JF. Identification of KIN (KIN17), a human gene encoding a nuclear DNA-binding protein, as a novel component of the TP53-independent response to ionizing radiation. Radiat Res. 2001;156: 535–544.

35. Biard DSF, Miccoli L, Despras E, Harper F, Pichard E, Créminon C, et al. Participation of kin17 protein in replication factories and in other DNA transactions mediated by high molecular weight nuclear complexes. Mol Cancer Res. 2003;1: 519–531.

36. Maga G, Biard DSF, Angulo JF. The human stress-activated protein kin17 belongs to the multiprotein DNA replication complex and associates in vivo with mammalian replication origins. and cellular biology. 2005. Available: https://mcb.asm.org/content/25/9/3814.short

37. Angulo JF, Rouer E, Mazin A, Mattei MG, Tissier A, Horellou P, et al. Identification and expression of the cDNA of KIN17, a zinc-finger gene located on mouse chromosome 2, encoding a new DNA-binding protein. Nucleic Acids Res. 1991;19: 5117–5123.

38. Gao X, Liu Z, Zhong M, Wu K, Zhang Y, Wang H, et al. Knockdown of DNA/RNA‑binding protein KIN17 promotes apoptosis of triple‑negative breast cancer cells. Oncology Letters. 2018. doi:10.3892/ol.2018.9597

39. Zhang Y, Huang S, Gao H, Wu K, Ouyang X, Zhu Z, et al. Upregulation of KIN17 is associated with non-small cell lung cancer invasiveness. Oncology Letters. 2017. pp. 2274–2280. doi:10.3892/ol.2017.5707

40. Valens M, Bohn C, Daignan-Fornier B, Dang VD, Bolotin-Fukuhara M. The sequence of a 54.7 kb fragment of yeast chromosome XV reveals the presence of two tRNAs and 24 new open reading frames. Yeast. 1997;13: 379–390.

41. Biard DSF, Miccoli L, Despras E, Frobert Y, Creminon C, Angulo JF. Ionizing radiation triggers chromatin-bound kin17 complex formation in human cells. J Biol Chem. 2002;277: 19156–19165.

42. Tran NT, Taverna M, Miccoli L, Angulo JF. Poly(ethylene oxide) facilitates the characterization of an affinity between strongly basic proteins with DNA by affinity capillary electrophoresis. Electrophoresis. 2005;26: 3105–3112.

43. Timchenko T, Bailone A, Devoret R. Btcd, a mouse protein that binds to curved DNA, can substitute in Escherichia coli for H-NS, a bacterial nucleoid protein. EMBO J. 1996;15: 3986–3992.

44. Miccoli L, Biard DSF, Créminon C, Angulo JF. Human kin17 protein directly interacts with the simian virus 40 large T antigen and inhibits DNA replication. Cancer Res. 2002;62: 5425–5435.

45. Cloutier P, Lavallée-Adam M, Faubert D, Blanchette M, Coulombe B. Methylation of the DNA/RNA-binding protein Kin17 by METTL22 affects its association with chromatin. J Proteomics. 2014;100: 115–124.

46. Mazin A, Milot E, Devoret R, Chartrand P. KIN17, a mouse nuclear protein, binds to bent DNA fragments that are found at illegitimate recombination junctions in mammalian cells. Mol Gen Genet. 1994;244: 435–438.

47. Tissier A, Kannouche P, Mauffrey P, Allemand I, Frelat G, Devoret R, et al. Molecular cloning and characterization of the mouse Kin17 gene coding for a Zn-finger protein that preferentially recognizes bent DNA. Genomics. 1996;38: 238–242.

48. Pinon-Lataillade G, Masson C, Bernardino-Sgherri J, Henriot V, Mauffrey P, Frobert Y, et al. KIN17 encodes an RNA-binding protein and is expressed during mouse spermatogenesis. J Cell Sci. 2004;117: 3691–3702.

49. le Maire A, Schiltz M, Braud S, Gondry M, Charbonnier J-B, Zinn-Justin S, et al. Crystallization and halide phasing of the C-terminal domain of human KIN17. Acta Crystallogr Sect F Struct Biol Cryst Commun. 2006;62: 245–248.

50. Miccoli L, Biard DSF, Frouin I, Harper F, Maga G, Angulo JF. Selective interactions of human kin17 and RPA proteins with chromatin and the nuclear matrix in a DNA damage- and cell cycle-regulated manner. Nucleic Acids Res. 2003;31: 4162–4175.

51. Le MX, Haddad D, Ling AK, Li C, So CC, Chopra A, et al. Kin17 facilitates multiple double-strand break repair pathways that govern B cell class switching. Sci Rep. 2016;6: 37215.

52. Kannouche P, Mauffrey P, Pinon-Lataillade G, Mattei MG, Sarasin A, Daya-Grosjean L, et al. Molecular cloning and characterization of the human KIN17 cDNA encoding a component of the UVC response that is conserved among metazoans. Carcinogenesis. 2000;21: 1701–1710.

53. Rappsilber J, Ryder U, Lamond AI, Mann M. Large-scale proteomic analysis of the human spliceosome. Genome Res. 2002;12: 1231–1245.

54. Makarov EM, Makarova OV, Urlaub H, Gentzel M, Will CL, Wilm M, et al. Small nuclear ribonucleoprotein remodeling during catalytic activation of the spliceosome. Science. 2002;298: 2205–2208.

55. Herold N, Will CL, Wolf E, Kastner B, Urlaub H, Lührmann R. Conservation of the protein composition and electron microscopy structure of Drosophila melanogaster and human spliceosomal complexes. Mol Cell Biol. 2009;29: 281–301.

56. Sidhar SK, Clark J, Gill S, Hamoudi R, Crew AJ, Gwilliam R, et al. The t(X;1)(p11.2;q21.2) translocation in papillary renal cell carcinoma fuses a novel gene PRCC to the TFE3 transcription factor gene. Hum Mol Genet. 1996;5: 1333–1338.

57. Skalsky YM, Ajuh PM, Parker C, Lamond AI, Goodwin G, Cooper CS. PRCC, the commonest TFE3 fusion partner in papillary renal carcinoma is associated with pre-mRNA splicing factors. Oncogene. 2001;20: 178–187.

58. Weterman MA, van Groningen JJ, Tertoolen L, van Kessel AG. Impairment of MAD2B-PRCC interaction in mitotic checkpoint defective t(X;1)-positive renal cell carcinomas. Proc Natl Acad Sci U S A. 2001;98: 13808–13813.

59. Agafonov DE, Deckert J, Wolf E, Odenwälder P, Bessonov S, Will CL, et al. Semiquantitative proteomic analysis of the human spliceosome via a novel two-dimensional gel electrophoresis method. Mol Cell Biol. 2011;31: 2667–2682.

60. Hegele A, Kamburov A, Grossmann A, Sourlis C, Wowro S, Weimann M, et al. Dynamic protein-protein interaction wiring of the human spliceosome. Mol Cell. 2012;45: 567–580.

61. Au V, Li-Leger E, Raymant G, Flibotte S, Chen G, Martin K, et al. CRISPR/Cas9 Methodology for the Generation of Knockout Deletions in Caenorhabditis elegans. G3. 2019;9: 135–144.

62. Hodgkin J, Papp A, Pulak R, Ambros V, Anderson P. A new kind of informational suppression in the nematode Caenorhabditis elegans. Genetics. 1989;123: 301–313.

63. Mitrovich QM, Anderson P. Unproductively spliced ribosomal protein mRNAs are natural targets of mRNA surveillance in C. elegans. Genes Dev. 2000;14: 2173–2184.

64. Kent WJ, Sugnet CW, Furey TS, Roskin KM, Pringle TH, Zahler AM, et al. The Human Genome Browser at UCSC. Genome Research. 2002. pp. 996–1006. doi:10.1101/gr.229102

65. Chanarat S, Sträßer K. Splicing and beyond: the many faces of the Prp19 complex. Biochim Biophys Acta. 2013;1833: 2126–2134.

66. Erkelenz S, Poschmann G, Ptok J, Müller L, Schaal H. Profiling of cis- and trans-acting factors supporting noncanonical splice site activation. RNA Biology. 2021. pp. 118–130. doi:10.1080/15476286.2020.1798111

67. Tarn WY, Steitz JA. SR proteins can compensate for the loss of U1 snRNP functions in vitro. Genes Dev. 1994;8: 2704–2717.

68. Brenner S. The genetics of Caenorhabditis elegans. Genetics. 1974;77: 71–94.

69. Wicks SR, Yeh RT, Gish WR, Waterston RH, Plasterk RH. Rapid gene mapping in Caenorhabditis elegans using a high density polymorphism map. Nat Genet. 2001;28: 160–164.

70. Dobin A, Davis CA, Schlesinger F, Drenkow J, Zaleski C, Jha S, et al. STAR: ultrafast universal RNA-seq aligner. Bioinformatics. 2013;29: 15–21.

71. McKenna A, Hanna M, Banks E, Sivachenko A, Cibulskis K, Kernytsky A, et al. The Genome Analysis Toolkit: a MapReduce framework for analyzing next-generation DNA sequencing data. Genome Res. 2010;20: 1297–1303.

72. Cingolani P, Platts A, Wang LL, Coon M, Nguyen T, Wang L, et al. A program for annotating and predicting the effects of single nucleotide polymorphisms, SnpEff: SNPs in the genome of Drosophila melanogaster strain w1118; iso-2; iso-3. Fly. 2012;6: 80–92.

73. Cvitkovic I, Jurica MS. Spliceosome database: a tool for tracking components of the spliceosome. Nucleic Acids Res. 2013;41: D132–41.

74. Doudna JA, Charpentier E. The new frontier of genome engineering with CRISPR-Cas9. Science. 2014. Available: https://science.sciencemag.org/content/346/6213/1258096.abstract?casa_token=OrPPcX2ZwwkAAAAA:cEKODhc7qG22k1LWzJyk_aCF7ZoU4eyQFxEqzbtWZ9P0xBpIDP6RhelPzEwBv8ybpJ7WFC-lz57C

75. Haeussler M, Schönig K, Eckert H, Eschstruth A, Mianné J, Renaud J-B, et al. Evaluation of off-target and on-target scoring algorithms and integration into the guide RNA selection tool CRISPOR. Genome Biol. 2016;17: 148.

76. Paix A, Folkmann A, Rasoloson D, Seydoux G. High Efficiency, Homology-Directed Genome Editing in Caenorhabditis elegans Using CRISPR-Cas9 Ribonucleoprotein Complexes. Genetics. 2015;201: 47–54.

77. Arribere JA, Bell RT, Fu BXH, Artiles KL, Hartman PS, Fire AZ. Efficient marker-free recovery of custom genetic modifications with CRISPR/Cas9 in Caenorhabditis elegans. Genetics. 2014;198: 837–846.

78. Crooks GE, Hon G, Chandonia J-M, Brenner SE. WebLogo: a sequence logo generator. Genome Res. 2004;14: 1188–1190.

79. Madeira F, Park YM, Lee J, Buso N, Gur T, Madhusoodanan N, et al. The EMBL-EBI search and sequence analysis tools APIs in 2019. Nucleic Acids Res. 2019;47: W636–W641.

